# Decomposing multi-scale dynamic regulation from single-cell multiomics with scMagnify

**DOI:** 10.64898/2026.02.03.703669

**Authors:** Xufeng Chen, Xi Yan, Bihan Shen, Heqi Wang, Zhixuan Tang, Yan Zang, Ping Lin, Haibing Zhang, Yu Li, Hong Li

**Author notes:** Corresponding author Hong Li, Yu Li.

## Abstract

Deciphering the highly coupled regulatory circuits that drive cellular dynamics remains a fundamental goal in biology. However, capturing the multi-scale time-lagged dynamics and combinatorial regulatory logic of gene regulation remains computationally challenging. Here we present scMagnify, a deep-learning-based framework that leverages multiomic single-cell assays of chromatin accessibility and gene expression via nonlinear Granger causality to reconstruct and decompose multi-scale gene regulatory networks (GRNs). Benchmarking on both simulated and real datasets demonstrates that scMagnify achieves superior performance. scMagnify employs tensor decomposition to systematically identify combinatorial TF modules and their activation profiles across different time-lags. It enables a hierarchical dissection of the regulatory landscape— from the activity of individual regulator to the combinatorial logic of regulatory modules and intercellular communications. We applied scMagnify to human hematopoiesis and mouse pancreas development, where it successfully recovered known lineage-driving regulators and provided novel insights into the combinatorial logic that governs cell fate decisions. Furthermore, in the context of kidney injury, scMagnify’s intracellular communication module mapped key signaling-to-transcription cascades linking microenvironment cues to pathological epithelial cell changes. In summary, scMagnify provides a powerful and versatile computational framework for dissecting the multi-scale regulatory logic that governs complex biological processes in development and disease.

## Introduction

Cell fates and complex organismal structures are established through a sophisticated series of molecular cues that orchestrate intricate and highly specific programs of gene expression. These programs are governed by interconnected molecular networks, which are shaped by a repertoire of cell-type-specific transcription factors (TFs)^1–3^. Importantly, TFs typically bind to cis-regulatory regions containing clusters of diverse TF binding sites. When associated TFs are co-expressed in overlapping spatial domains, their combinatorial binding generates precise spatiotemporal transcriptional activity patterns^4,5^. This context-dependent regulation allows TFs to integrate multiple signals, activating or repressing target genes (TGs) with high specificity, thereby driving the intricate gene expression programs required for cellular specialization and organismal development^6^.

Over the past decades, a variety of computational methods have been proposed to unravel the intricate regulatory mechanisms controlling gene expression^7,8^. Transcriptome-based methods such as WGCNA^9^, GENIE3^10^, GRNBoost2^11^ and PIDC^12^ infer TF–TG relationships by capturing covariation in gene expression. However, these methods produce undirected networks, cannot distinguish direct from indirect effects, and lack causal interpretability^13,14^.

To overcome these limitations, recent advances have taken two major directions. First, GRN inference methods that incorporate (pseudo)temporal information leverage the temporal ordering of cells to align the expression patterns of candidate TFs with those of their putative TGs, often using time lag-based correlations. This temporal resolution improves inference accuracy and enables more causal interpretations of regulatory relationships^15–18^, a crucial limitation remains. Their mathematical foundations require a sequential ordering of cells, which struggles to handle complex cell-state landscapes with branching or convergent fates. Velorama^19^ overcomes this limitation by using partial ordering-based Granger causality to model non-linear cellular dynamics. Second, additional data modalities such as chromatin accessibility (e.g., DNase-seq^20^, ATAC-seq^21^) or TF binding profiles (e.g., ChIP–seq^22^) have been integrated to provide mechanistic evidence. These allow the identification of regulatory elements (REs), which can be linked to TFs via motif matching and associated with nearby TGs, forming TF–RE–TG connections^23^. While these strategies offer mechanistic insights, they often rely on separate measurements from different samples or cell populations, limiting their resolution and ability to resolve dynamic regulatory relationships.

The development of single-cell sequencing has begun to overcome these challenges by profiling the transcriptome and epigenome at cellular resolution and enabled the investigation of causal and temporal relationships among critical regulatory events^24–26^. Pioneering methods like SCENIC^27^ reconstructed cell-type-specific GRNs by integrating scRNA-seq data with population-level chromatin information. Subsequent approaches, including Pando^28^, FigR^29^, and SCENIC+^30^, introduced various optimizations by leveraging single-cell chromatin accessibility data in more intricate ways to better define regulatory connections. Furthermore, LINGER^31^ employs a lifelong learning framework to build highly accurate GRNs by integrating knowledge from atlas-scale external bulk data across diverse cellular contexts. Beyond improving GRN inference accuracy, a new generation of methods has introduced powerful downstream applications to provide more biological meaningful insights. For example, CellOracle^32^ utilizes the inferred GRNs to simulate the effects of TF perturbations and predict their function in driving cell state transitions, while Dictys^33^ moves beyond a static view by modeling how the GRNs dynamically rewire across a continuous cellular process.

Despite these significant advances, several key limitations remain^26^. First, for inference accuracy, most methods do not simultaneously leverage multimodal and pseudotemporal information, which are critical for identifying causal regulatory interactions. Second, for downstream application, current approaches tend to focus on individual regulator activity in discrete cell states, overlooking the continuous dynamics of cellular transitions and the combinatorial logic^5^ of regulatory modules central to complex biological processes.

Here, we present scMagnify (**M**ulti sc**A**le **G**ene regulatory **N**etwork **I**n**F**erence and anal**Y**sis), an integrated computational framework designed to address these challenges. To improve inference accuracy, scMagnify synergistically integrates single-cell multi-omic data and pseudotemporal information through a nonlinear Granger causality model^34,35^ to infer multi-scale GRNs. For downstream analysis, it enables a multi-level investigation of gene regulation, moving from individual regulators to the identification of combinatorial regulatory modules via tensor decomposition^36^, and further linking these intracellular dynamics to extracellular cues by modeling signaling-to-transcription cascades. We demonstrate the power and versatility of scMagnify in both developmental and disease contexts, where it rediscovers known master regulators and reveals novel, multi-scale regulatory programs driving cell state transitions.

## Results

### Overview of scMagnify

scMagnify is a versatile Python toolkit for inferring and analyzing gene regulatory networks (GRNs) from single-cell multi-omic datasets. To ensure a targeted and robust GRN inference, scMagnify first identifies a set of high-confidence genes intrinsically linked to the specified cellular dynamics. It achieves this by establishing cellular trajectories and using Generalized Additive Models (GAMs)^37^ to select genes whose expression significantly correlates with the pseudotime axis. To further constrain the model with plausible chromatin interactions, scMagnify then constructs a basal TF binding network for each cell state. This structural scaffold is built by linking genes to potential cis-regulatory elements (cCREs) via expression-accessibility correlation and scanning these regions for known TF binding motifs^38^. Together, these steps yield a focused, biologically relevant search space for the final GRN inference (**Fig. 1a and Methods**).

**Fig. 1.**
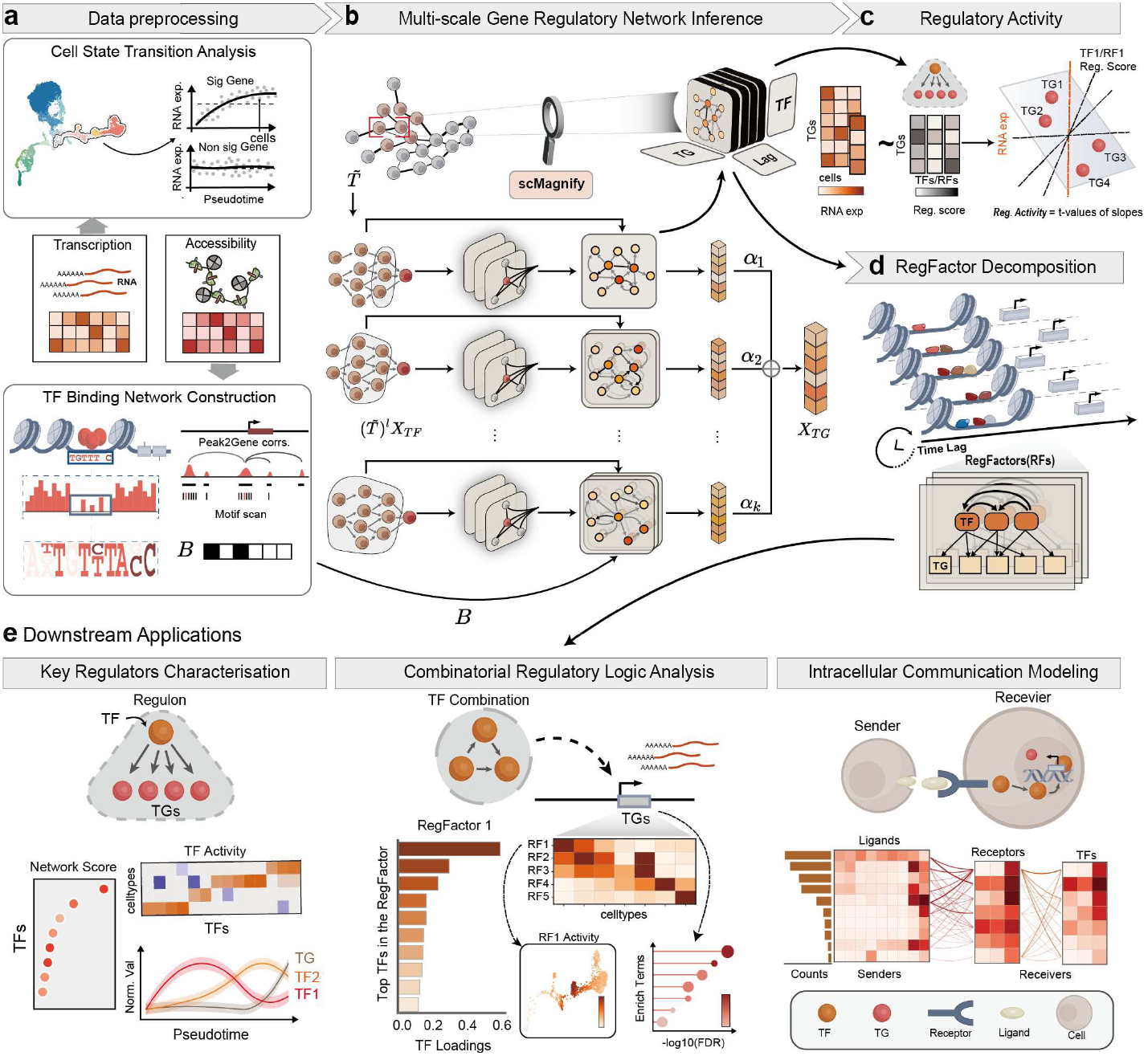
Overview of the scMagnify framework. **a, Data preprocessing**. scMagnify integrates single-cell transcriptomic and chromatin accessibility data. It first identifies genes with dynamic expression along a pseudotime trajectory. Concurrently, it constructs a basal GRN by linking transcription factors (TFs) to target genes (TGs) based on peak-to-gene correlations and TF motif scanning. **b, Multi-scale GRN inference**. A self-explaining neural network implementing nonlinear Granger causality infers the GRN. The model uses a multi-scale architecture with parallel branches to capture regulatory effects across different time lags, constrained by the basal GRN. An attention mechanism integrates these signals to predict TG expression. **c, Regulatory activity inference**. The activity of a TF or RF is quantified using a multivariate linear model based on the expression of its target regulon. The resulting t-value from the model fit serves as the final activity score. **d, RegFactor decomposition**. The inferred multi-scale regulatory scores are organized into a third-order tensor (TFs, TGs, time-lags). Tucker decomposition is applied to this tensor to deconvolve co-regulating TF modules, termed RegFactors (RFs). **e, Downstream applications**. The inferred GRNs and RegFactors enable multiple analyses: (left) characterization of key regulators by network centrality and activity dynamics over pseudotime; (middle) analysis of regulatory synergy through TF loadings, activity patterns, and functional enrichment of RegFactors; and (right) modeling of intercellular communication by linking ligand-receptor pairs to downstream TF activation in receiver cells.

Within this focused search space from the previous step, scMagnify performs GRN inference using a partial ordering-based^19,39^ Granger causality framework. This framework is implemented via a self-explaining neural network featuring a multi-scale architecture. More specifically, we consider these lagged TF expression values as predictors for the regulation coefficient matrices in a feedforward neural network. To capture regulatory effects across different time scales, the model uses parallel branches, each examining cellular dynamics over a different retrospective time lag. Finally, an attention mechanism then automatically learns to weight the information from each branch, integrating these multi-scale signals to construct the final GRN (**Fig. 1b and Supplementary Fig. 1 and Methods**).

scMagnify includes a suite of functions to analyse context-specific GRNs. To isolate the most robust interactions, the network is filtered based on user-defined statistical thresholds (**Methods**). The resulting GRN then serves as a basis for functional and structural analysis. A primary application is the inference of regulatory activity, where the activity of each TF is quantified from the collective expression of its TGs (its regulon) using a multivariate linear model^40^ (**Fig. 1d and Methods**). Beyond this functional insight, the network structure can be investigated through standard graph-theoretic metrics to identify pivotal hub regulators within the cellular circuitry (**Fig. 1e**).

It is well-established that TFs often act in a combinatorial manner to regulate their target genes^41,42^. To systematically explore these regulatory synergies within dynamic processes, scMagnify employs a modified Tucker decomposition approach. This regfactor decomposition method is applied to a tensor comprising the regulatory coefficients (Reg. scores) derived from the multi-scale Granger causality model. This approach enables the simultaneous identification of co-regulating TF modules and their shared sets of TGs across different time lags (**Fig. 1c, e and Supplementary Fig. 1b and Methods**).

Furthermore, scMagnify links intercellular communication to intracellular regulation by dissecting signaling-to-transcription cascades along the pseudotime axis. To achieve this, it first identifies key pathways consisting of ligand-receptor pairs and their downstream TFs from prior knowledges^43–45^. It then correlates each receptor’s expression with its target TF’s expression across pseudotime and validates these interactions with a permutation test^46^. This analysis reveals robust signaling axes that are active during specific cellular transitions (**Fig. 1e and Methods**).

Collectively, scMagnify dissects the multi-level architecture of gene regulation—from the activity of individual regulators, to the combinatorial logic of regulatory modules, and finally to the systemic coordination via intercellular communication—enabling mechanistic insights into the regulation of cellular dynamics.

### scMagnify demonstrates superior performance across extensive benchmarks

To provide a comprehensive evaluation, we benchmarked scMagnify against a set of leading GRN inference methods (**Supplementary Table1**). The selected methods represent a variety of modeling strategies and differ in their core capabilities. For example, a major class of methods including LINGER, Dictys, CellOracle and FigR excel at integrating multi-omic data to infer GRNs, but they do so in a static manner without explicitly modeling temporal dynamics. Conversely, another class of methods, such as Velorama^19^ and SINCERITIES^16^, can model cellular dynamics using pseudotime information, but they are limited to single-cell transcriptomic data. This benchmark design allows us to specifically highlight the advantages of scMagnify’s integrated framework, which uniquely combines multi-modal data with pseudotemporal information to reconstruct GRNs. We also reported the results from scMagnify without the TF binding network (scMagnify-nobasal) to isolate the contribution of the chromatin prior and to enable a fair comparison with methods that rely solely on transcriptomic data.

Our evaluation was performed on both synthetic datasets with known ground-truth network structures and well-characterized real-world multi-omic datasets. We systematically compared the methods from multiple perspectives, including their accuracy in predicting TF-TG interactions, their ability in recovering key driver TFs, and the computational stability and scalability (**Fig. 2a**).

**Fig. 2.**
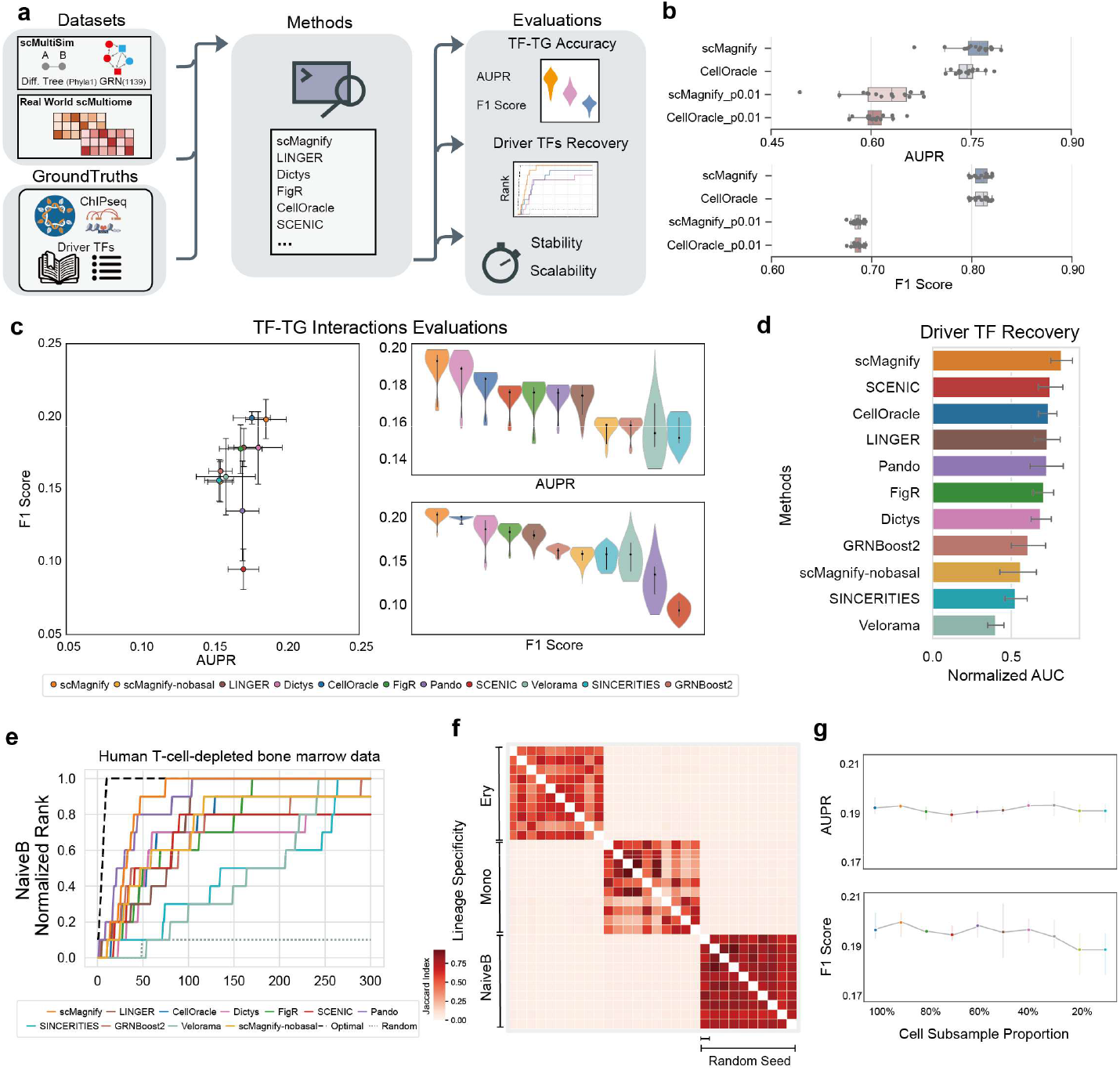
scMagnify demonstrates superior performance in comprehensive benchmarks. **a**, Schematic of the benchmarking strategy. **b**, Performance on synthetic datasets. Boxplots show the AUPR and F1-scores for scMagnify and CellOracle on data with standard and high noise levels. **c**, Evaluation of TF-TG interaction inference on real-world data. (Left) A scatter plot compares the mean AUPR and F1-score for each method across nine hematopoietic lineages. Error bars represent the standard error of the mean. (Right) Violin plots show the distribution of AUPR and F1-scores for each method. **d**, Driver TF recovery benchmark. A bar plot shows the normalized AUC for the recovery of known lineage-driving TFs, aggregated across all lineages. Error bars indicate the standard error of the mean. **e**, Rank recovery curves show the cumulative fraction of known driver TFs identified within the top-ranked TFs for the Naive B cell lineage. **f**, Evaluation of reproducibility and lineage specificity. The heatmap displays the Jaccard similarity between GRNs inferred from 10 independent runs (using different random seeds) across three distinct lineages. **g**, Robustness to cell subsampling. Line plots show AUPR and F1-scores (y-axis) plotted against the fraction of the cell population used (x-axis).

First, we applied scMultiSim^47^ to generate synthetic single-cell multiomics datasets with different parameters (**Supplementary Fig. 2a**). We used the area under the precision-recall curve (AUPR) and the F1-score to assess the accuracy of ground-truth network reconstruction, as both metrics are well-suited for the sparse nature of GRNs.The primary finding is that scMagnify achieves a superior median AUPR in both standard and noisy settings (p0.01)^47^ (Methods), underscoring its overall high accuracy. While its performance profile shows a broader variance with noisy data, the central tendency of its predictions remains consistently higher than CellOracle’s (**Fig. 2b**).

Next, to evaluate scMagnify’s real-world performance, we applied our cell state transition module to uniformly processed single-cell multi-omic datasets, identifying 9 distinct cellular lineages for subsequent benchmark (**Supplementary Fig. 2b-g and Methods**). Each of these lineages represents a unique dynamic process and served as an independent test set for GRN inference. Then we curated a corresponding gold-standard set of TF-TG interactions from a comprehensive ChIP-seq database to serve as ground truth. Specifically, we collected 1016 ChIP-seq experiments for 234 TFs from the Cistrome database^48^ to assess whether each method could successfully recover these experimentally-supported TF-TG interactions, with performance quantified by the AUPR and F1-score (**Methods**). Despite performance variations across methods and lineages, the collective evidence points to the superior overall performance of scMagnify. Specifically, an analysis of the full performance distributions reveals that scMagnify consistently obtained the highest median AUPR and F1-score (**Fig. 2c, d and Supplementary Fig. 3a-c**). This indicates that its typical performance is robustly superior to all competitors. In conclusion, our systematic set of benchmarks on real-world data demonstrates the comparable or superior performance of scMagnify, highlighting its reliability and high accuracy across a range of dynamic cellular processes.

While edge-level accuracy is a fundamental metric for GRN inference, a more stringent test of any inferred GRN is whether its network topology can be used to identify biologically important regulators^49,50^. We therefore performed a driver TF recovery benchmark to assess if scMagnify could prioritize known lineage-driving TFs (**Supplementary Table 2**) more effectively than competing methods. For each inferred network, we ranked TFs by their degree centrality and measured how well this ranking recovered a curated gold-standard list of drivers for each lineage. As illustrated by the rank recovery curves (**Fig. 2d, e and Supplementary Fig. 4a, b**), the ranking produced by scMagnify showed the strongest enrichment for known drivers. An aggregate score, summarizing the overall recovery performance, further confirmed this trend, with scMagnify ranking first among all tested methods (**Fig. 2d**). This suggests that the topology of the GRN from scMagnify is not only accurate but also structurally meaningful, successfully identifying key regulators based on their network connectivity.

A critical test for any GRN inference method is its ability to generate networks that are both reproducible and context-specific^33^. To assess scMagnify against this standard, we compared GRNs generated from 10 independent runs with different random seeds across 3 distinct lineages. The resulting Jaccard similarity heatmap revealed a clear block-diagonal structure, a pattern arising from high intra-lineage similarity (reproducibility) and low inter-lineage similarity (specificity) (**Fig. 2f**). Moreover, the method proved highly robust to cell subsampling, with both AUPR and F1 scores remaining stable even when the cell population was reduced to 20% (**Fig. 2g**). These results validate that scMagnify can robustly infer GRNs that are both consistent within a given biological context and unique to it.

Comprehensive benchmarking demonstrates scMagnify’s superior performance in GRN inference, where its novel multi-omic integration framework enables highly accurate, robust, and context-specific reconstruction of regulatory networks and identification of key TFs.

### scMagnify dissects lineage-specific key regulators in human hematopoiesis

To demonstrate the utility of scMagnify in developmental contexts, we obtained a human blood dataset containing single-cell multi-omic data of CD34+ bone marrow cells from healthy donors^51^, a paradigmatic system where tightly regulated transcriptional programs drive the differentiation of hematopoietic stem cells into a complex hierarchy of distinct lineages^52^. We first classified the data into three lineages and selected 3,281 genes that were significantly associated with pseudotime (FDR < 0.0001, A > 0.3) (**Supplementary Fig. 2b-d and Methods**). As shown in Fig. 3a, HSCs follow 3 main developmental trajectories—the erythroid, monocyte, and Naïve B cell lineages— which are defined by a progression through distinct intermediate cell states. Then we applied scMagnify to investigate the lineages separately to construct the lineage-specific GRN and identify the key regulators driving cell fate decisions.

**Fig. 3.**
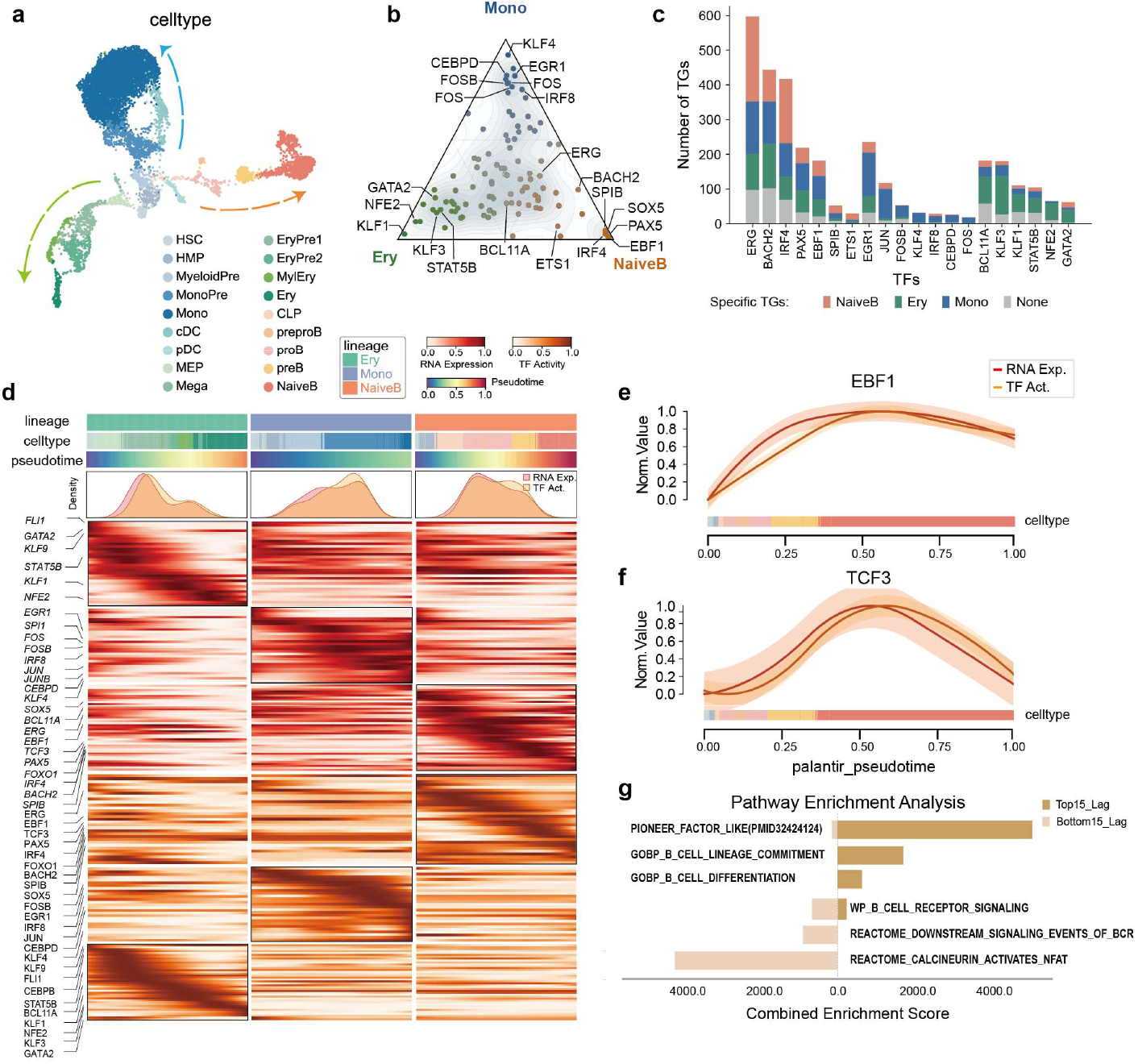
scMagnify dissects lineage-specific regulatory programs in human hematopoiesis. **a**, UMAP embedding of CD34+ bone marrow cells, colored by cell type. Arrows indicate the three major differentiation trajectories: erythroid, monocyte, and Naïve B cell. **b**, Ternary plot of key TFs associated with the erythroid, monocyte, and Naive B cell lineages. TFs are positioned according to their lineage specificity. **c**, Bar plot quantifying the number and lineage specificity of target genes (TGs) for key TFs. Bar colors represent targets specific to Naive B, erythroid, or monocyte lineages, or those shared across lineages. **d**, Heatmap displaying the dynamics of TF gene expression (top panel) and inferred regulatory activity (bottom panel) along the pseudotime axis for the three hematopoietic lineages. **e, f**, Plots showing the temporal decoupling between normalized RNA expression (orange line) and inferred TF activity (purple line) for B-cell regulators EBF1 (**e**) and TCF3 (**f**) along the B-cell lineage pseudotime. Shaded areas represent the 95% confidence interval. **g**, Pathway enrichment analysis comparing TFs with the longest regulatory lags (Top15_Lag) and the shortest regulatory lags (Bottom15_Lag). The x-axis shows the combined enrichment score. HSC, Hematopoietic Stem Cell;HMP, Hematopoietic Multipotent Progenitor;CLP, Common Lymphoid Progenitor;MEP, Megakaryocyte-Erythroid Progenitor;MyeloidPre, Myeloid Progenitor; MonoPre, Monocyte Progenitor;Mono, Monocyte;cDC, Conventional Dendritic Cell;pDC, Plasmacytoid Dendritic Cell;Mega, Megakaryocyte;Ery, Erythroid;EryPre1, Erythroid Progenitor 1; EryPre2,Erythroid Progenitor 2;MylEry, Myeloid-Erythroid Progenitor;preproB, Pre-pro-B cell; proB, Pro-B cell;preB, Pre-B cell;NaiveB, Naive B-cell.

By comparing GRNs across different lineages (**Methods**), scMagnify identified major lineage-defining TFs with highly specific target sets, including EBF1, PAX5, IRF4 and SPIB for the B cell lineage^53–56^; GATA2 and KLF1 for the erythroid lineage^57,58^; and IRF8, EGR1 and KLF4 for the monocyte lineage^59^ (**Fig. 3b, c and Supplementary Fig. 5a, b**). In contrast, the analysis also identified TFs with broader, non-lineage-specific regulatory roles. For instance, ERG^60^ and BACH2^61^ regulate a large number of targets that are distributed across two or more lineages, indicating their function as more general regulators in hematopoiesis (**Fig. 3c**).

To better understand how transcriptional programs are orchestrated in different lineages, we next analysed the transcriptional and regulatory dynamics along trajectories. This analysis revealed distinct waves of TF expression that define each lineage, with key regulatory modules being activated in a highly specific temporal order (**Fig. 3d**). Specifically, the erythroid trajectory is characterized by the strong early expression of a module including GATA2 and KLF1, the monocytic (Mono) trajectory is driven by the subsequent induction of pioneer factors such as CEBPD and CEBPB, while the Naive B-cell lineage is established through the robust co-activation of key B-cell determinants like EBF1, PAX5, and BCL11A ^62^(**Supplementary Table 2)**.

Our analysis also highlights the temporal decoupling between the expression of a transcription factor and its regulatory activity^30,63^ (**Fig. 3d and Methods**). This is exemplified by the B-cell determinants EBF1 and TCF3^64^, whose expression precedes the full activation of the downstream gene program required for lineage commitment (**Fig. 3e, f)**. By inferring GRNs that operate across multiple time-lag scales, scMagnify provides a framework to quantify the regulatory lag of individual transcription factors (**Methods**).

Consistent with previous studies, we found that this regulatory lag is a key indicator of a TF’s biological role^19,65^. TFs with a longer lag tend to be major hubs in the regulatory network, showing a strong positive correlation with degree centrality (**Supplementary Fig. 5c-e**). Functionally, these long-lag TFs, which include the B-cell determinants EBF1, PAX5, and TCF3, are highly enriched for foundational processes like “pioneer-like properties^66^”, “B cell differentiation,” and “Cell fate commitment.” In contrast, TFs with shorter lags are associated with more immediate cellular responses, such as “B cell receptor signaling” and “BCR signaling events” (**Fig. 3g)**.

In summary, scMagnify systematically dissects the key regulators and their dynamic regulation across different hematopoietic lineages. These regulators are characterized by distinct regulatory lags, highlighting their temporal roles as either foundational hubs or rapid-response effectors. Building on this principle of temporal regulation, we next investigated the combinatorial regulatory logic governing B cell commitment.

### scMagnify recovers regulatory logic governing B cell fate commitment

Deciphering the regulatory logic that drives B cell commitment is challenging, as the process relies on temporally orchestrated combinatorial inputs rather than a simple linear TF cascade^67–69^. To systematically explore these dynamic regulatory patterns, we therefore applied scMagnify’s regfactor decomposition, hypothesizing that distinct co-regulating TF modules controlling distinct cellular programs can be deconvolved from our multi-scale GRNs.

By decomposing the multi-scale GRNs, we identified several RegFactors. Analysis revealed that each of these RegFactors is regulated by a different combination of TFs, activated at a different developmental stage, and is associated with distinct biological functions.

Specifically, our analysis identified 5 RegFactors that orchestrate B-cell lineage (**Fig. 4a and Supplementary Fig.6a-c**). RegFactor 1, predominantly active in the earliest progenitor states (HSC/HMP), includes key TFs such as MECOM, RORA, and PBX1, suggesting its role in stem cell maintenance, proliferation and alternative lineage priming^70–72^. From this stage, progenitors in HMP/CLP initiate lineage priming through the activation of RegFactor 2, which serves as a key transitional program. This is followed by RegFactor 3, which is highly enriched in cell-cycle regulators^73,74^ like E2F3 and TFDP1 and drives the robust proliferation of these primed progenitors, preparing them for subsequent lineage commitment. In contrast, cells that commit to the B-lineage downregulate these programs and instead activate RegFactor 4 during the pro-B and pre-B stages. Governed by master regulators IRF4, PAX5, TCF3 and EBF1, these factors are central to establishing B-cell identity^64^. Finally, RegFactor 5, driven by ERG, IRF4, governs the functional readiness of B cells by orchestrating the molecular machinery required for both antigen recognition and presentation^75^. (**Fig. 4b, c and Supplementary Table 3**).

**Fig. 4.**
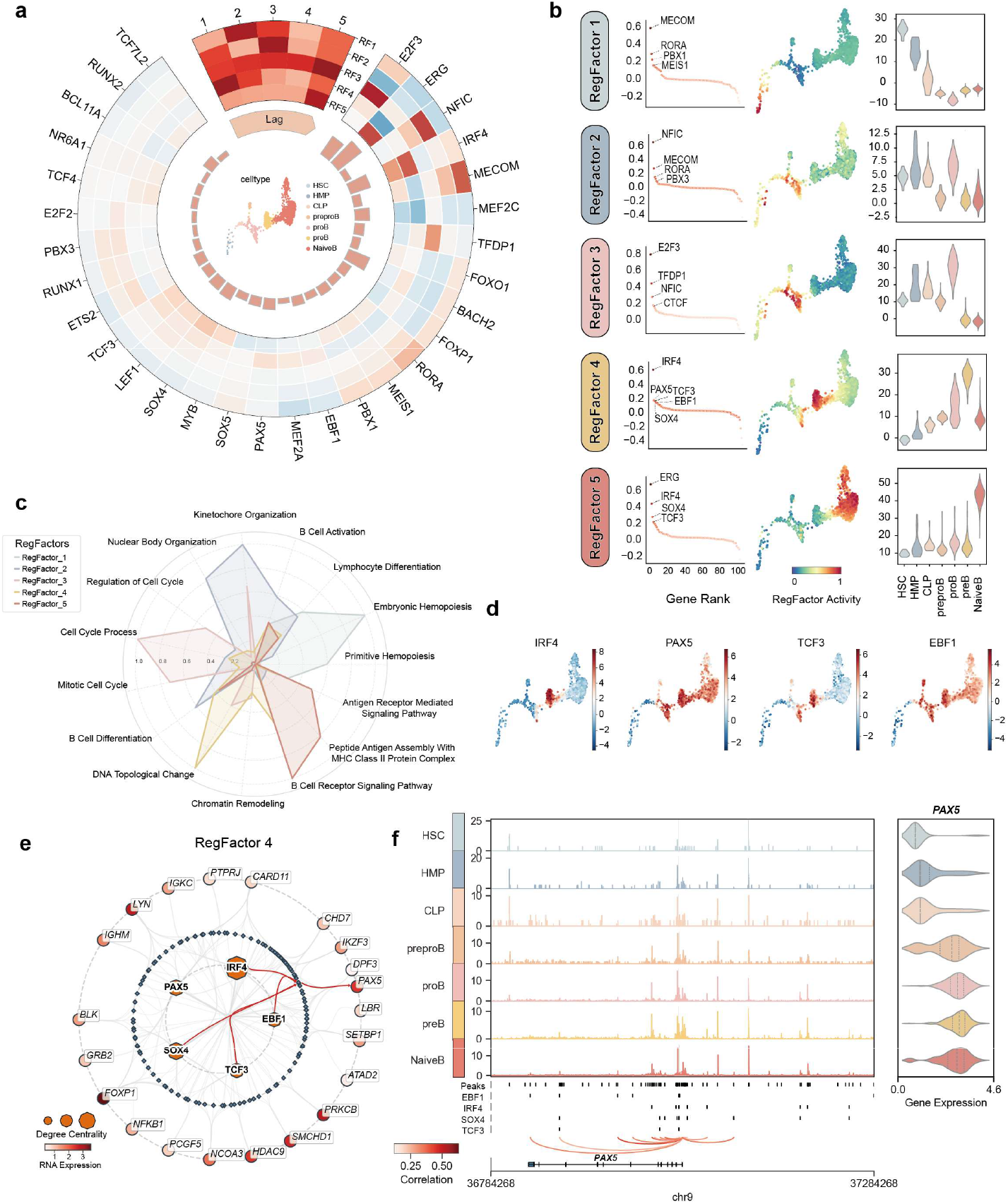
RegFactor decomposition reveals combinatorial regulatory logic driving B cell commitment. **a**, A circular heatmap summarizing the 5 RegFactors (RFs) identified in the Naive B cell lineage. From outer to inner rings, the plot shows contributing TFs, TF loadings for each RF, and the average regulatory lag of each RF. **b**, Characterization of the RegFactors. For each RF, the plots show the top TFs by TF loading (left bar plot), the activity projected on the UMAP embedding (middle), and the activity distribution across cell types (right violin plot). **c**, A radar plot showing the results of functional enrichment analysis for each RegFactor. Each axis represents a distinct biological pathway. **d**, Trajectory plots showing the inferred regulatory activity of the four core TFs of RegFactor 4 (IRF4, PAX5, TCF3, and EBF1) along the B-cell differentiation path. **e**, A subnetwork view of RegFactor 4, organized in three layers to illustrate the regulatory cascade. Inner hexagonal nodes represent the core TFs. The middle ring of diamond nodes represents candidate cis-regulatory elements (cCREs). The outer ring of circular nodes represents target genes (TGs). For TFs, node size corresponds to degree centrality and for TGs, node color indicates scaled RNA expression. Red edges highlight the specific regulatory interactions targeting the PAX5 gene, corresponding to the genomic analysis in panel f. **f**, Genomic view of the PAX5 locus. Chromatin accessibility tracks are shown for different stages of B cell development. The bottom tracks indicate the positions of cCREs (peaks) and predicted binding motifs for key RegFactor 4 TFs. Violin plots on the right show PAX5 RNA expression across the corresponding cell types.

To understand the molecular programs underlying B cell commitment, we focused our investigation on RegFactor 4. Consistent with the overall activity of RegFactor 4, its key TFs showed tightly-correlated activity patterns peaking in the pro-B and pre-B cell stages (**Fig. 4d and Supplementary Fig.6d**). Together, these core TFs orchestrate a broad downstream program of TGs critical for B-cell function, including components for immunoglobulin chains (*IGHM, IGKC*), B-cell receptor signaling (*BLK, LYN*)^76^, and chromatin modification (*CHD7, HDAC9*)^77^ (**Fig. 4e and Supplementary Fig. 6**).

To further assess scMagnify’s capacity for inferring regulatory synergy, we examined PAX5, a master regulator of B-cell commitment and identity. Epigenetic analysis of the *PAX5* locus confirmed its co-regulation, showing its expression correlates with accessibility at distal enhancers linked to its promoter. Critically, these linked sites showed clear motif co-binding signatures for key RegFactor 4 TFs, including IRF4, EBF1, TCF3 and SOX4, providing direct epigenetic evidence for their co-regulatory role (**Fig. 4f**).

### scMagnify deciphers regulatory programs driving pancreatic lineage bifurcation

To test the versatility of scMagnify, we applied it to a single-cell multi-omic dataset of mouse embryonic pancreas development. This dataset encompasses the diversification of endocrine progenitors into the primary alpha (alpha) and beta (beta) cell fates, alongside delta (δ) and epsilon (ε) lineages (**Fig. 5a and Supplementary Fig. 6e, f**). We leveraged this developmental map to systematically identify the key TFs and their time-varying interactions that orchestrate alpha versus beta lineage bifurcation.

**Fig. 5.**
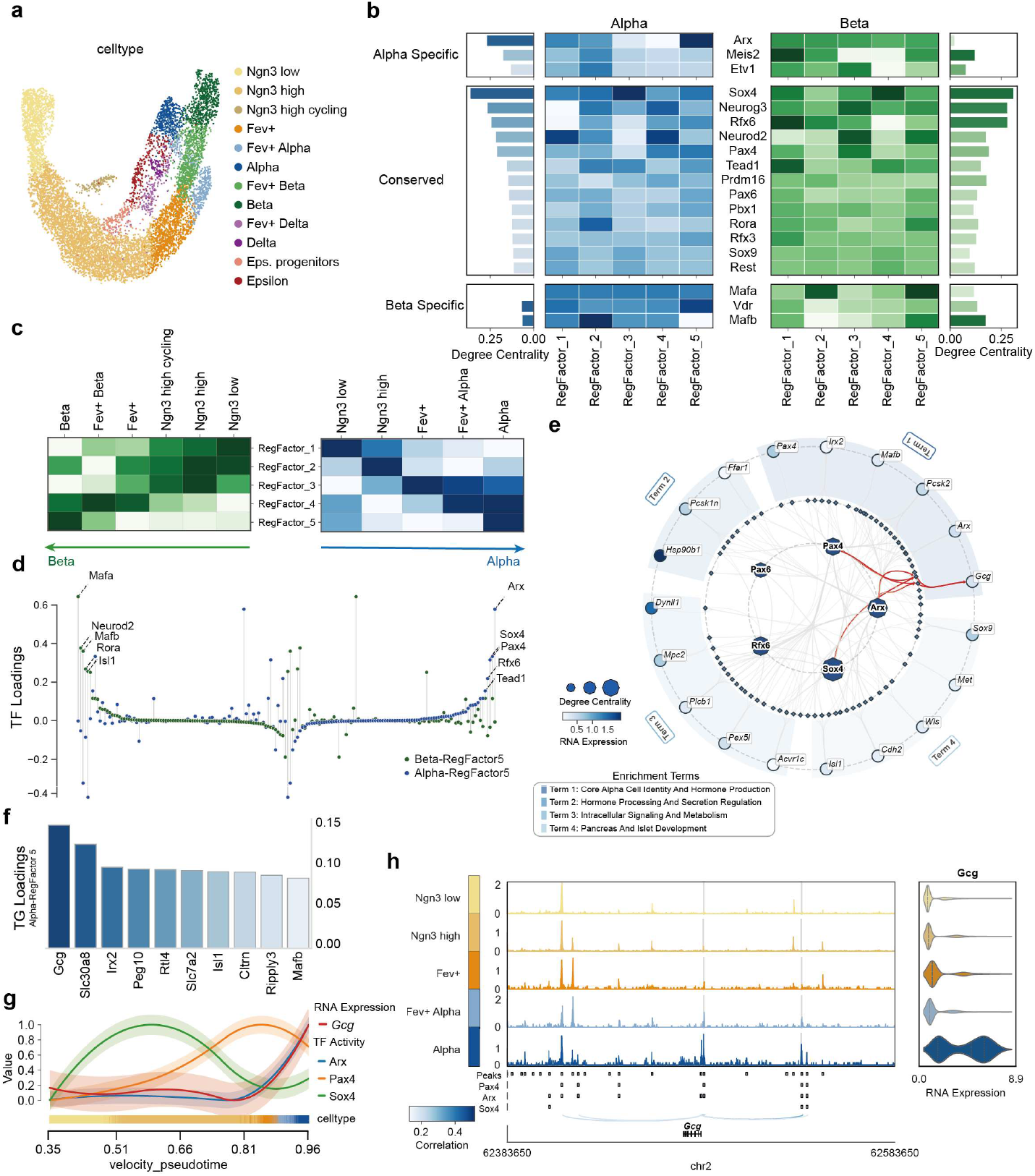
scMagnify dissects the regulatory programs driving pancreatic alpha and beta cell lineage bifurcation. **a**, UMAP embedding of mouse embryonic pancreas cells, colored by cell type, showing the endocrine differentiation landscape. **b**, Heatmaps showing TF contributions to the five RegFactors for the alpha (left) and beta (right) lineages. TFs are grouped as lineage-specific or conserved. Adjacent bar plots show the degree centrality for each TF. **c**, Heatmaps of the inferred activity for each of the five RegFactors across different cell states in the beta (left) and alpha (right) differentiation trajectories. **d**, Scatter plot of TF loadings for the terminal selector modules, Beta-RegFactor 5 and Alpha-RegFactor 5. **e**, Subnetwork view of the Alpha-RegFactor 5 module. Node size represents degree centrality and color indicates RNA expression. Functionally related target genes are grouped by enrichment terms. **f**, Bar plot of the top target gene (TG) loadings for the Alpha-RegFactor 5 module. **g**, Line plot showing the dynamics of Glucagon (Gcg) gene expression and the inferred regulatory activity of its key TFs (Arx, Pax4, Sox4) along the alpha-cell pseudotime trajectory. **h**, Genomic view of the Gcg locus. Tracks show chromatin accessibility across different cell states of the alpha lineage. Bottom tracks indicate predicted binding motifs for key TFs. Violin plots on the right show Gcg gene expression across the corresponding cell types. Ngn3 low/high/high cycling, Endocrine progenitors with low, high, or high cycling expression of Neurog3; Fev+, FEV-positive progenitor; Eps. progenitors, Epsilon cell progenitors; Alpha, Alpha cell; Beta, Beta cell; Delta, Delta cell; Epsilon, Epsilon cell.

Based on this dataset, scMagnify inferred dynamic, multi-scale GRNs that capture the regulatory shifts during cell fate decision. It revealed a cohort of lineage-defining TFs whose regulatory activities peaked or shifted dramatically around the bifurcation point, underscoring their critical role in cell fate commitment (**Supplementary Fig. 7**).

To systematically identify the regulatory modules driving lineage specification, we employed the RegFactor decomposition. The GRNs deconvolved into 5 distinct RegFactors for each lineage, revealing 3 classes of TFs: alpha-specific, beta-specific, and a larger group of conserved regulators active in both lineages (**Fig. 5b and Supplementary Table 4, 5**).

For the alpha cell lineage, Arx emerged as the key hub regulator, supported by Meis2 and Etv1, consistent with its established role in defining alpha-cell identity^78–80^. Conversely, Mafa and Mafb were identified as key beta-specific TFs, consistent with their known roles in beta-cell function and maturation^81^. Alongside these lineage-specifiers, a conserved module of TFs—including Sox, Neurog3, and Rfx6—exhibited high centrality in both lineages, representing a core endocrine regulatory program^82–85^.

We next sought to characterize the temporal dynamics of these RegFactors. By projecting their activities along the pseudotime axis, we found that each factor displayed a unique activity profile along both the alpha and beta cell lineages (**Fig. 5c and Supplementary Fig 7 a, b**). This revealed that the lineage-specific RegFactor 5 for each fate exhibited the most striking stage-specific pattern. In both lineages, the activity of their respective RegFactor 5 surged dramatically only at the final maturation stage, identifying these distinct TF sets as the “terminal selectors”^86^ that lock in final cell identity. To identify the specific TFs driving these terminal programs, we examined the TF loadings for each lineage’s RegFactor 5 (**Fig. 5d**). The analysis confirmed they are composed of largely distinct TF cohorts: the alpha-specific module is overwhelmingly dominated by Arx^87^, while the beta-specific module is principally led by Mafa^88^. This quantitatively pinpoints the distinct master regulators executing the final fate decision in each lineage.

To gain a mechanistic view of how the alpha-specific terminal module establishes cell identity, we visualized the Alpha-RegFactor 5 regulatory network (**Fig. 5e**). This revealed a sophisticated architecture organized around a core of co-regulating TFs, with Arx acting as the unequivocal central hub. Its dense connections to other module members, such as Pax4 and Sox4, highlight the combinatorial nature of this regulatory program.

Most critically, the network provides a direct mechanistic link between this module and the quintessential function of an alpha cell. We identified a strong, direct regulatory connection from Arx to *Gcg* (Glucagon), the definitive hormone produced by alpha cells (**Fig. 5g**). Underscoring this critical link, an analysis of TG loadings showed that *Gcg* is the highest-weighted gene in this module (Fig. 5f). Beyond direct hormone synthesis, the module coordinates the broader functional machinery required for its activity. For example, it targets the prohormone convertase *Pcsk2*, essential for processing proglucagon, and other genes involved in secretion modulation like *Ffar1*^89^. Epigenetic analysis confirms this link, showing *Gcg* expression correlates with accessibility at enhancers co-bound by key module TFs, including Arx, Pax4, and Sox4 (**Fig. 5h**).

### scMagnify maps signaling-to-transcription cascades driving epithelial cell adaptation during kidney injury

Intercellular communication via ligand-receptor interactions is fundamental to coordinating cellular responses in development and disease. However, how these extracellular signals are dynamically translated into specific intracellular transcriptional programs to orchestrate cell state transition remains largely unresolved^46,90^. To address this, we developed a dynamic communication module within scMagnify that links intercellular signaling to intracellular TF activity along a pseudotemporal trajectory. Briefly, after establishing ligand-receptor-TF links from a knowledgebase, we aggregated cells into metacells ordered along a pseudotime trajectory. We then correlated receptor and TF expression across these metacells and employed a permutation test to identify robust, time-varying signaling-to-transcription axes. (**Fig. 6a and Methods**).

**Fig. 6.**
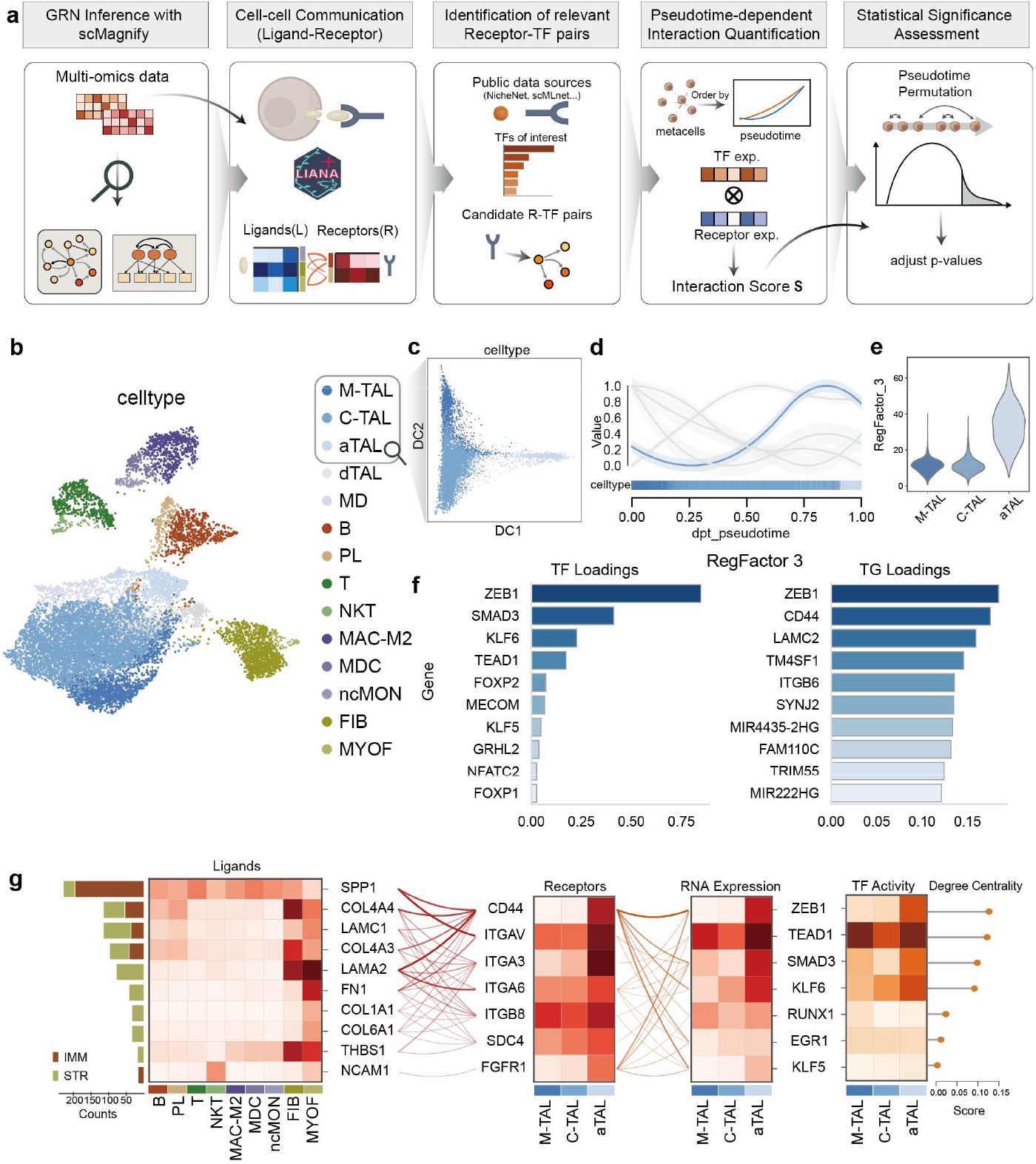
scMagnify maps signaling-to-transcription cascades driving epithelial cell adaptation during kidney injury. **a**, A schematic overview of the scMagnify intracellular communication module. The workflow integrates GRN inference, ligand-receptor analysis, and pseudotime ordering to quantify and statistically assess signaling-to-transcription cascades. **b**, UMAP embedding of single cells from the human kidney cortex, colored by cell type. **c**, A diffusion map of the thick ascending limb (TAL) epithelial cell populations, showing the inferred injury trajectory from healthy (M-TAL, C-TAL) to adaptive (aTAL) states. **d**, A line plot showing the inferred activity of RegFactor 3 along the TAL pseudotime trajectory. **e**, A violin plot comparing the activity of RegFactor 3 across the M-TAL, C-TAL, and aTAL cell states. **f**, Bar plots showing the TF loadings (left) and TG loadings (right) for the aTAL-specific RegFactor 3. **g**, A summary plot of the key signaling-to-transcription cascades targeting RegFactor 3 TFs. Heatmaps show the expression of ligands in various sender cell populations (left), the expression of corresponding receptors in TAL cells (middle), and the RNA expression and activity of downstream TFs in TAL cells (right). Lines connect ligands to their receptors and receptors to downstream TFs. M-TAL, Medullary Thick Ascending Limb; C-TAL, Cortical Thick Ascending Limb ; aTAL, Adaptive Thick Ascending Limb; dTAL, Distal Thick Ascending Limb; MD, Macula Densa; B, B cell; PL, Plasma cell; T, T cell; NKT, Natural Killer T cell; MAC-M2, M2-like Macrophage; MDC, Monocyte-derived Dendritic Cell; ncMON, Non-classical Monocyte; FIB, Fibroblast; MYOF, Myofibroblast; IMM, Immune cells; STR, Stromal cells.

To validate scMagnify’s capability to infer signaling pairs driving pathological cell state transitions in a complex disease setting, we analyzed a single-cell multi-omic dataset from the cortex of patients with acute and/or chronic kidney disease (AKI/CKD)^91^ (**Fig. 6b**). While this dataset comprises diverse kidney cell types, our analysis focuses on the epithelial populations to investigate their adaptive responses to injury. The dataset captures the transition of thick ascending limb (TAL) epithelial cells from a healthy, cortical state (C-TAL) to a dysregulated adaptive state (aTAL). Trajectory inference using a diffusion map resolved a continuous progression from healthy (C-TAL/M-TAL) to adaptive TAL cell states (aTAL), establishing a pseudotemporal axis to dissect the dynamics of the injury response (**Fig. 6c and Supplementary Fig. 8b**).

Building on the inferred trajectory, we constructed the GRNs for the TAL, allowing us to identify key regulators **(Supplementary Fig. 8a)**, and then decomposed the network into 5 RegFactors **(Fig. 6d, Supplementary Fig. 8d-f and Supplementary Table 6)**. Notably, RegFactor 3, which governs TGs functionally enriched for pathological processes like cell adhesion and signaling (**Supplementary Fig. 8c**), exhibited activity that was highly and specifically enriched in the aTAL state (**Fig. 6d, e**). It is principally driven by the pro-fibrotic TFs ZEB1, SMAD3 and KLF6^92–95^, which are upregulated in the aTAL state (**Fig. 6f**). This prompted us to focus our analysis on this module to dissect the regulatory program driving the adaptive injury response.

To identify the key upstream signaling axes, we prioritized ligand-receptor pairs based on the correlation strength between the receptor’s expression and the expression of the core RegFactor 3 TFs (**Fig. 6f, g**). The top-ranked interaction was the SPP1-CD44/ITGAV axis. Computationally, SPP1 was the most abundant ligand and originated from a broad set of microenvironment cells, including both stromal (STR) and immune (IMM) populations. Its receptor, CD44, was significantly upregulated on aTAL cells, creating a potent pro-fibrotic signaling channel^96^. In contrast, our analysis also identified the NCAM1-FGFR1 signaling axis. Unlike the widespread sourcing of SPP1, the NCAM1 signal originated almost exclusively from immune cells, highlighting a highly specific mode of communication that connects to downstream TFs such as EGR1 and RUNX1 (**Fig. 6g**). These findings demonstrate scMagnify’s ability to deconvolve the complex signaling landscape into both broad-acting and cell-type-specific pathological pathways.

## Discussion

In this manuscript, we present scMagnify, an integrated computational framework designed to reconstruct, decompose, and analyze dynamic, multi-scale gene regulatory networks (GRNs) from single-cell multi-omic datasets. To achieve this, scMagnify synergistically integrates single-cell multi-omic data with pseudotemporal information through a nonlinear Granger causality model implemented within an interpretable multi-scale neural network. The inference is further constrained by chromatin accessibility-derived priors to enhance accuracy. A key innovation of our framework is its ability to reconstruct the GRN as a multi-scale tensor, explicitly capturing time-lagged regulatory interactions along a dynamic process. This unique structure is pivotal, as it enables subsequent analyses, such as tensor decomposition to identify dynamic regulatory modules and the linking of these modules to upstream signaling-to-transcription cascades.

Our comprehensive benchmarks on both synthetic and real-world datasets demonstrate that scMagnify achieves state-of-the-art performance, consistently outperforming leading methods in both network reconstruction and the critical task of identifying driver TFs. Beyond benchmarks, the application of scMagnify to complex biological systems yielded significant novel insights. In the context of development, our analysis of human hematopoiesis and mouse pancreas development revealed the temporal decoupling of transcription factor expression from its regulatory activity and identified the specific terminal selector modules that orchestrate cell fate bifurcation. In a disease context, scMagnify successfully mapped the signaling cascades from the microenvironment that drive pathological cell state transitions in kidney injury. These findings, which are often missed by conventional analyses, underscore the power of a multi-scale approach.

While scMagnify represents a significant advance, we acknowledge several limitations and areas for future development. The framework’s performance is dependent on the quality of the input multi-omic data and the accuracy of the upstream trajectory inference. Furthermore, the construction of the basal GRN, while effective, relies on peak-to-gene correlations. Future iterations could incorporate more sophisticated approaches, such as transcription factor footprinting or advanced peak-to-gene linking strategies, to define these chromatin-based priors with even higher confidence. Another important consideration is that scMagnify, like other methods based on Granger causality, uses a predictive model for causal inference. Although powerful for identifying putative regulatory drivers, it does not capture underlying biochemical kinetics in the same way as mechanistic models based on principles like RNA velocity^97,98^. Consequently, achieving in silico perturbation and counterfactual inference remains a significant challenge that represents an important direction for future work^99^. The current Granger causality model is also built on a directed acyclic graph structure, which may not fully capture biological processes involving complex, cyclical dynamics^100^. Finally, while the current framework focuses on integrating transcriptome and chromatin accessibility data, future iterations could be extended to incorporate additional data modalities such as single-cell proteomics^101^ or ^102^ to provide an even more comprehensive view of gene regulation.

The advent of single-cell multi-omics is providing an unprecedented opportunity to understand complex biological processes with high resolution. To translate this rich data into a mechanistic understanding, it is essential to move beyond static, single-scale views of gene regulation. Frameworks like scMagnify, which embrace the dynamic, multi-scale, and combinatorial nature of regulatory networks, are critical for this endeavor. By providing a tool to dissect these complex systems, scMagnify can help generate novel, testable hypotheses and will be a valuable resource for advancing our understanding of development and disease. Ultimately, integrating such dynamic regulatory maps will be an essential step toward realizing the goals of large-scale initiatives aiming to build “virtual cells,” which seek to create predictive, multi-scale computational models of cellular function and behavior^103,104^.

## Methods

scMagnify is a computational framework to infer gene regulatory networks (GRNs) and explore dynamic regulation synergy from single-cell multiomics data. The construction of GRNs is predominantly achieved through nonlinear multivariable Granger causal inference, using an interpretable multi-scale neural network. Moreover, scMagnify integrates prior knowledge of chromatin conformation by incorporating a domain-specific constraint into model structure.

The scMagnify workflow consists of several steps: (1) Multi-omic data preprocessing (2) Multi scale gene regulatory network inference. (3) Downstream regulation analysis.

We implemented and tested scMagnify in Python and designed it for use in the Jupyter notebook environment. scMagnify code is open source and available on GitHub (https://github) along with detailed descriptions of functions and tutorials.

### Multi-omic Data Preprocessing

#### Cell state transition analysis

Cell state transition can be modeled as a multi-scale process. Existing approaches such as CellRank combines similarity-based markov models with multiview single-cell datas for cellular fate mapping.

##### Transition matrix construction

We begin by applying CellRank to calculate cell–cell transition probabilities, represented by a transition matrix *T* ∈ ℝ^*N×N*^, which reflects the likelihood of one cell transitioning to another along the pseudotime trajectory, which we index by *t*.

##### Cell state selection

Cells are then assigned to specific fates or lineages through clustering based on their fate probabilities. This step groups cells that are likely to follow similar developmental paths, allowing for the identification of distinct lineages.

##### Feature association test

Feature association is tested using generalized additive models (GAM), which model gene expression as a function of pseudotime *t* with cubic spline regression. The GAM model is compared to an unconstrained model using an F-test, and p-values are adjusted for multiple testing with an FDR threshold. This identifies key genes associated with cell state transitions along pseudotime.

#### TF binding network construction

For each cell state, scMagnify constructs a TF binding network that contains unweighted, directional edges between a TF and its TG. scMagnify uses the peak-gene correlations and TF-binding motifs for this task.

##### Identification of cCREs via peak-gene correlations

To identify candidate cis-regulatory elements (CREs), scMagnify first aggregates the single-cell multiomics data into metacells using SEACells^51^. For each gene, the method then computes the Pearson correlation between its normalized metacell expression and the normalized accessibility of each ATAC peak located within 100 kb upstream and 100 kb downstream. The significance of each peak-gene correlation is assessed against an empirical null distribution, which is generated by sampling 100 background peaks matched for GC content and accessibility^105^. This analysis identifies the subset of regions significantly correlated with gene expression, retaining pairs that meet the criteria of a correlation coefficient >0.1 and an empirical P-value < 0.1.

##### Motif scanning

Following the identification of cCREs, we scanned their DNA sequences to identify TF binding motifs, thereby linking TFs to potential TGs. This motif scanning was performed for all candidate regions using the MOODS^106^. By default, scMagnify uses the HOCOMOCO v11 core collection^107^ for human and mouse as its motif database. The results were aggregated into a binary adjacency matrix, hereafter referred to as the basal TF binding network ***B*** ∈ {**0, 1**}^***P***×***Q***^. An entry *B*_*p,q*_ = 1 indicates that a motif for *TF*_*p*_ was found within a cCRE linked to*TG*_*q*_, establishing a potential regulatory edge. In addition to this network, scMagnify also reports the specific genomic coordinates of each identified TF-binding site.

### Multi-scale Gene Regulatory Network Inference

To address the inherent limitations of classical Granger causal inference in handling dynamical systems with branching points, scMagnify employs a directed acyclic graph (DAG)-structured partial ordering^19^ to establish robust temporal relationships among variables. This approach ensures a principled representation of causal dependencies in complex systems, particularly in single-cell genomics.

#### Input data and preprocessing

The training of scMagnify relies on three primary inputs:

1. A gene expression matrix *X*, where *X* ∈ ℝ^*N×M*^, with *N* cells and *M* genes.
2. A cell transition matrix *T*, where *T* ∈ ℝ^*N×N*^, encoding pairwise relationships between cells.
3. A TF binding network *B* ∈ {**0, 1**}^***P***×***Q***^, which incorporates prior biological knowledge to guide the inference process.

##### Preprocessing of *X*

To ensure data comparability, raw expression counts were first normalized by scaling each cell’s library size to the median library size of the dataset. The resulting normalized counts were then log-transformed to stabilize variance and reduce skewness.

##### Preprocessing of *T*

To account for variability in the in-degrees of the DAG, we precomputed a modified matrix 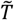, derived from the normalized transpose of *T* .Specifically, 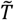 is defined such that each row sums to 1, ensuring a consistent scaling of influence across cells. For rows of *T*^*T*^ (the transpose of *T*) that consist entirely of zeros, no normalization is applied. Formally, 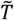 is computed as:

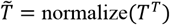

where the normalization ensures 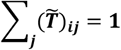 for all rows ***i*** with non-zero entries. This matrix 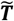 serves as a graph diffusion operator, enabling the aggregation of information over each observation’s ancestors in the DAG.

#### Model structure

Our model infers dynamic regulatory relationships by conceptualizing gene regulation as a sophisticated, non-linear extension of the classical Vector Autoregressive (VAR) framework, which is foundational to Granger causality^108^. The model’s architecture is built up progressively to address the complexities of biological systems. The foundational linear VAR model posits that TG expression can be predicted from a linear combination of past TF expression:

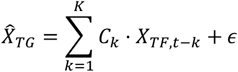

Here, *C*_*k*_ represents a constant coefficient matrix for a given time lag *k*. To capture the non-linear dynamics inherent in gene regulation, we first extend this model by replacing the constant matrix *C*_*k*_ with a neural network Ψ_*k*_ : ℝ^*P*^ → ℝ^*P×Q*^, which learns a multi-scale regulatory coefficient matrix based on historical TF expression^34^. To ensure biological plausibility, we impose a structural constraint on these learned coefficients by applying an element-wise multiplication with a prior TF binding network, *B*. This constrained non-linear model is formulated as:

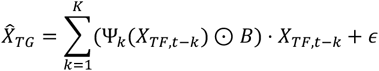

The final model further advances this framework to simultaneously account for regulatory effects across multiple time scales and to integrate the cell-state transition topology (**Supplementary Fig. 1a**). This is achieved through a multi-branch architecture with an attention mechanism, which operates on features propagated along a cell transition graph. The complete model can be formulated as:

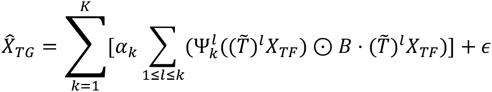

In this architecture, the model is composed of *K* independent and parallel branches, where each branch *k* acts as a “receptive field” considering historical information up to *k* steps away (**Supplementary Fig. 1b**). Instead of simple time lags, we use a graph diffusion operator 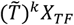, to aggregate TF expression information from *k*-hop ancestral cells in the trajectory graph, thereby embedding cell-state relationships into the temporal model.

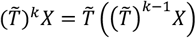

Within each branch *k*, an additional index *l* (where 1≤ *l* < *k*) is used to sum over all intermediate time lags, allowing the model to capture both immediate and delayed causal effects. Finally, an additional neural network produces learnable attention weights *α*_*k*_, which determine the emphasis placed on each branch, effectively merging all extracted causal information based on its relevance. Here, 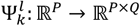 is a neural network parameterized by *θ*_*k*_ that includes an input layer, fully connected hidden layers with ReLU activation, and an output layer of size *P* × *Q*. (**Supplementary Fig. 1c**). The model’s direct output is the multi-scale regulatory coefficient tensor 𝒯, which quantifies the dynamic, context-specific regulatory strengths as determined by the attention-weighted aggregation of the neural network outputs:

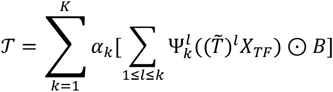

#### Training procedure

To mitigate spurious inference in multivariate time series, scMagnify is trained by minimizing the following penalized loss function using mini-batch gradient descent:

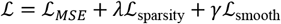

where *λ* and *γ* are hyperparameters that control the strength of the regularization terms. The components of the loss function are defined as:

#### Mean Squared Error (MSE) ℒ_*MSE*_

This term quantifies the difference between the model’s predicted target gene expression 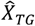, and the true expression values *X*_*TG*_ . It is calculated as the standard mean squared error:

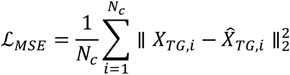

where *N*_*e*_ is the number of cells in the batch.

#### Sparsity-Inducing Regularization ℒ_sparsity_

To promote a sparse GRN and select only the most salient regulatory interactions, we employ a Group Elastic Net penalty^109^ on the outputted regulatory coefficient tensor 𝒯 . This penalty encourages the model to either drive all coefficients for a given TF-TG pair across all time lags to zero (group sparsity via L2 norm) or to selectively zero out individual time-lag-specific coefficients (element-wise sparsity via L1 norm). Let ***τ***_*p,q*_ *= (*𝒯_,*p,q*_, …, 𝒯_*K,p,q*_) be the vector of regulatory coefficients for TF *p* and TG *q* across all *K* time lags. The regularization term is defined as:

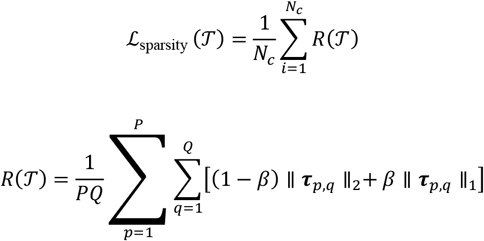

where *P* and *Q* are the number of TFs and TGs, respectively, and *β* is a hyperparameter that mixes the L1 and L2 penalties.

#### Temporal Smoothness Regularization ℒ_smooth_

To ensure that the inferred regulatory relationships evolve smoothly over the pseudotime trajectory, we introduce a penalty on the difference between the coefficient tensors of temporally adjacent cells. This term penalizes large, abrupt changes in the GRN structure between consecutive time points. The smoothness loss is calculated as the squared L2 norm of the difference between the coefficient tensor at pseudotime *t*_*i*_ and the tensor at the next time point *t*_*i*+1_:

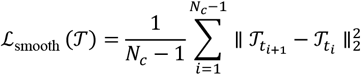

#### Model inference and postprocessing

After training, the model is applied to the entire dataset to infer cell-specific regulatory dynamics and subsequently post-processed to generate formatted results for downstream analysis (**Supplementary Fig. 1d**). For each of the *N* cells in the cell state, the model outputs a regulatory coefficient tensor. These individual tensors are first collected into a comprehensive tensor, which we denote as 𝒯_total_ ∈ ℝ^*N×K×P×Q*^ . This tensor represents the complete set of inferred regulatory coefficients across all cells, time lags, TFs, and TGs.

To derive a robust, context-level GRN, we perform a series of post-processing and aggregation steps. First, to ensure the reliability of inferred interactions, we filter out regulatory coefficients that exhibit high variability across the cell population. This is achieved by calculating the coefficient of variation (CV) for each interaction across the cell dimension of 𝒯_total_. Interactions with a CV below 0.2 are retained, resulting in a filtered tensor 𝒯_final_.

From this stabilized tensor, we construct the final GRNs. To obtain a representative multi-scale network that preserves time-lag-specific information, we compute the median of 𝒯_final_. across the cell dimension. This yields the multi-scale network 𝒢_multi-scale_ ∈ ℝ^*K×P×Q*^

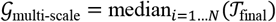

To create a single, consolidated GRN that summarizes the most dominant interactions across all time lags, we derive two ensemble networks. The ensemble network strength *G*_strength_, quantifies the maximum unsigned regulatory influence for each TF-TG pair and is constructed by taking the maximum value across the time-lag dimension (*k*) of the absolute median coefficients:

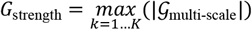

Similarly, the ensemble network activation identifies the dominant regulatory mode (activation or repression) by taking the maximum value of the signed median coefficients across the time-lag dimension:

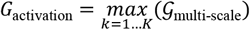

### Downstream Regulation Analysis

#### Network score calculation

The GRN can be investigated using general network topology metrics to identify key regulators. To facilitate this, we first perform network binarization. This step converts the continuous, weighted ensemble network strength matrix *G*_strength_, into a binary adjacency matrix by applying a user-defined sparsity threshold. On this resulting binary GRN, we then employ graph-theoretic approaches, such as calculating degree centrality, betweenness centrality, and PageRank, using the Python library networkx^110^ to characterize the network’s structure and pinpoint influential TFs.

#### Regulatory activity inference

Following this filtering step, we leverage the filtered GRN to infer transcription factor (TF) activities. Specifically, the activity of a TF is represented by the expression levels of its TGs, collectively referred to as a *regulon*.

To quantify the activity of each regulon, we used a multivariate linear model (MLM) using the decoupler^40^ package, which estimates regulator activities by jointly modeling all regulators in the network and assigning activity scores based on the t-values of the fitted model.

For each cell, the MLM estimates a vector of TF activities **a**_*TF*_, that best explains the TG expression vector **x**_*TG*_, given the ensemble network activation matrix *G*_activation_, which serves as the score matrix **W**:

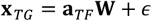

The final activity score for each TF is the t-value from the fitted model, which provides a normalized measure of activity.

#### RegFactor decomposition

To identify combinatorial regulatory modules, we apply Tucker decomposition to a third-order multi-scale signed network 𝒢_multi-scale_ ∈ ℝ^*K×P×Q*^ . This tensor is constructed by aggregating the regulatory coefficients, derived from the multi-scale Granger causality model, across the time-lags. The decomposition is formulated as:

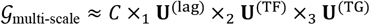

This formulation approximates the original three-way tensor 𝒢_multi-scale_, as the n-mode product of a smaller core tensor (*C*) and three factor matrices. The core tensor 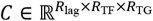, captures the interaction strengths between the decomposed modules. By default, the rank of the decomposition for each mode (*R*_lag_ × *R*_TF_ × *R*_TG_) is set to be equal to the number of time lags *K*. The three factor matrices define the modules themselves: the lag factor matrix 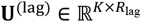 defines temporal activation profiles across the time lags; the TF factor matrix 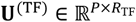 defines co-regulating TF modules; and the TG factor matrix 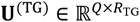 defines shared target gene modules. Together, a specific combination of a TF module, a TG module, and a temporal profile, linked by the core tensor, constitutes a ‘RegFactor’ and its activity can also be estimated using the MLM method, where the TG loadings from the corresponding column of the *U*^(TG)^ matrix serve as the weight matrix *W*.

#### Intracellular communication

To link intercellular communication with intracellular transcriptional regulation, scMagnify models signaling-to-transcription cascades along a pseudotime axis. The process begins by identifying ligand-receptor pairs and TFs of interest. Putative signaling pathways are then constructed by linking receptors to their downstream TFs using a prior knowledge base from scMLnetDB^44^.

For each candidate receptor-TF pair, an interaction score is quantified across the inferred trajectory. Cells are first aggregated into *M* metacells ordered by pseudotime to reduce noise. Let 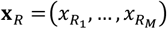 and 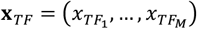 be the vectors of the receptor’s and TF’s average expression across the metacells, respectively. The interaction score is calculated as the sample covariance between these vectors:

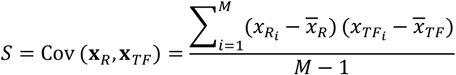

where 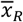 and 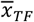 are the mean expression values.

The statistical significance of the observed score *S*_*obs*_, is assessed via a permutation test. The pseudotime ordering of the metacells is shuffled *N* times (by default, *N* = 1000), and a score 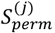, is calculated for each permutation *j* . This generates an empirical null distribution. The empirical p-value for a two-tailed test is then calculated as:

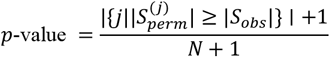

The resulting p-values are adjusted for multiple testing to identify significant signaling axes.

### Datasets

#### Simulated data

To benchmark the performance of existing GRN inference methods, we used scMultiSim to generate simulated single-cell multiomics datasets with the following configurations: 1139-gene GRN, 1,000 cells, 50 CIFs, rd = 0.2, σi = 1 and other default parameters. We generated a total of eight datasets with random seed from 1 to 8. Technical noise and batch effects were then added using default parameters. Specifically, the technical noise was modeled by randomly adding and removing connections in the simulated ATAC peak-gene matrices, where the probability of these modifications is controlled by a noise-level parameter *p*. ‘scale’ is set to 50 × *p*_*pos*_, where *p*_*pos*_ is the proportion of positive edges within each matrix.

#### Single-cell multi-omic data

Single-cell Multiome ATAC + Gene Expression data were obtained from previously published studies, including: (1) human hematopoiesis datasets from Persad et al., comprising CD34^+^-enriched and T cell-depleted bone marrow cells; (2) a mouse pancreas development dataset from Klein et al., consisting of embryonic pancreatic cells; and (3) human kidney disease datasets from Gisch et al., derived from percutaneous kidney biopsy samples.

The gene expression matrix (GEM) was normalized by total UMI counts and log-transformed using Scanpy with default parameters. The chromatin accessibility data had been deduplicated in the original studies. Cell annotations such as cell type and low-dimensional embeddings were also obtained from the original studies.

#### Human T-cell-depleted bone marrow data

The single-cell multi-omic data from Persad et al.^51^ were originally described and made available on Zenodo (https://doi.org/10.5281/zenodo.6383269). For this study, the preprocessed AnnData objects for gene expression (RNA) and chromatin accessibility (ATAC) were downloaded directly from Mellon^111^ tutorial. The cell type annotations (celltype) and Palantir-derived pseudotime values (palantir_pseudotime) were used directly from the metadata provided in these original files.

#### Human CD34+ bone marrow data

This dataset also originates from the study by Persad et al.^51^ (Zenodo: https://doi.org/10.5281/zenodo.6383269). For this study, the preprocessed AnnData objects for gene expression (RNA) and chromatin accessibility (ATAC) were downloaded directly from SEACells tutorial. Palantir-derived pseudotime values (palantir_pseudotime) were retained from the original data files. For this study, the cell type annotations (celltype) were re-annotated by markers.

#### Mouse pancreatic endocrine data

This dataset was originally published by Klein, D. et al.^112^, with raw data available from the Gene Expression Omnibus (GEO) under accession number GSE275562. The preprocessed MuData object (.h5mu), which contains the necessary AnnData objects for this study, was obtained as described in the mubind^113^ tutorial. RNA velocity-related data and the original celltype annotations were used directly from the metadata provided in this file.

#### Human kidney injury data

This dataset was originally published by Gisch et al.^91^ and made available on Zenodo (https://doi.org/10.5281/zenodo.8029990). The preprocessed Seurat objects were downloaded from this repository. The cell type annotations for this study were derived by merging and re-annotating the original subclass.l2 metadata. Pseudotime trajectories were computed using the Diffusion Pseudotime (DPT) algorithm^114^ from the Scanpy package^115^ (scanpy.tl.dpt).

### Evaluations

To evaluate whether scMagnify can correctly identify cell-state-specific GRN, we benchmarked our method against diverse GRN inference algorithms: LINGER, Dictys, CellOracle, FigR, Pando, SCENIC, Velorama and SINCERITITES.

#### Preparation of input data for GRN inference

For each of the two human bone marrow multi-omic datasets, we first performed feature association test to identify significantly genes (FDR < 0.001, A > 0.3). Next, we subsetted the cells into nine lineages (Ery/Mono/NaiveB for the T-cell-depleted dataset; pDC/cDC/Mono/Ery/CLP/Mega for the CD34+ dataset).

#### GRN inference methods

After preprocessing, the same dataset was used as input for each GRN inference algorithm to ensure a fair comparison. We followed the official tutorials for each package and used default hyperparameters unless stated otherwise. Notably, SCENIC+ was not included in this comparison, as its workflow depends on pycisTopic^116^ to perform cell type specific peak analysis. The benchmarking framework in this study is established on independent lineages, that is not directly compatible with the analytical steps required by pycisTopic.

#### Ground-truth data preparation

Cell-type-specific ground-truth GRNs were generated in the same manner as in a previous benchmarking study^33^. We obtained tissue-specific ChIP–seq peak data from the Cistrome database^48^. For each experiment (by GSMID), the top 100 peaks (by peak score) were regarded as positive hits and annotated to the nearest gene. The TF binding gold standard was defined as TF– target pairs that were positive hits in at least one of the experiments.

#### TF-TG interactions evaluation

To assess the accuracy of inferred TF-TG interactions, we benchmarked performance on both synthetic and real-world datasets. For both data types, each method’s ranked predictions were evaluated using the Area Under the Precision-Recall curve (AUPR) to assess overall ranking quality and the F1-score to evaluate the balance of precision and recall for a set of top-ranked predictions. The F1-score is calculated as:

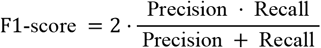

#### Driver TF recovery evaluation

To evaluate how well each method identifies key regulators, we performed a driver TF recovery benchmark^30,50^. First, for each inferred network, we ranked all TFs by their degree centrality *C*_*D*_, calculated as the sum of connections for each TF node i in the adjacency matrix A:

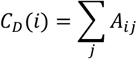

We then compared this ranking against a curated list of known driver TFs. To quantify performance, we generated a recovery curve, ϕ(N), which measures the number of known drivers found within the top N ranked TFs. The y-axis of this curve is the ‘Normalized Rank’, which represents the fraction of total known drivers recovered at rank N. The overall performance for each method m was summarized by calculating the Area Under this Curve (AUC):

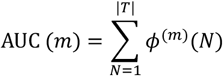

where ∣T∣ is the total number of TFs. A higher AUC score indicates a better performance in prioritizing known driver TFs.

#### Reproducibility and cell-type specificity evaluations

We evaluated network reproducibility and specificity across three distinct cell lineages. For each lineage, we generated 10 independent GRNs by running the inference algorithm with different random seeds.

The GRN similarity between any two inferred networks (A and B), defined by their sets of regulatory edges, was quantified using the Jaccard Index:

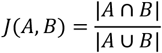

### Analysis

#### Functional enrichment analysis

Functional enrichment was conducted using Over-Representation Analysis (ORA) with gene sets from the Molecular Signatures Database (MSigDB). Statistical significance was assessed using a one-sided Fisher’s exact test, which computes the probability of enrichment based on the hypergeometric distribution.

The p-value for observing an overlap of size *o* (given a query list of size *r*, a gene set of size *g*, and a background universe of *D*) was calculated as:

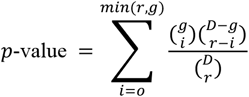

Enriched terms were ranked by a ‘Combined score’, integrating the p-value and the Odds Ratio (OR):

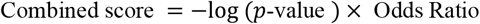

All resulting p-values were adjusted for multiple testing using the Benjamini-Hochberg FDR correction.

#### Regulation specificity analysis

To quantify regulation specificity, two matrices were compiled for each gene across the Ery, NaiveB, and Mono lineages: 1) the absolute mean TF activity and 2) the mean expression. Each matrix was independently normalized on a per-gene basis by dividing the gene’s value in each lineage by its total sum across all three lineages. A final composite specificity score was computed by averaging these two resulting proportional (normalized) matrices.

Lineage-specific TGs were determined by first calculating the log_2_(Fold Change) for each gene’s expression in one lineage against the average expression of the other two lineages. A gene was then assigned to a specific lineage if its maximum log_2_(FC) ≥0.4 and this maximum value was also at least 0.2 greater than its median log_2_(FC) across all three lineages.

#### Regulation lag calculation

scMagnify quantifies the characteristic regulatory lag for each transcription factor (TF) based on the inferred multi-scale network tensor, 𝒢_multi-scale_ ∈ ℝ^*K×P×Q*^.

First, for each TF-TG interaction, the vector of regulatory coefficients across all *K* lags is L1-normalized. This converts the coefficients into a distribution of relative regulatory influence across the different time lags. An estimated lag for the specific interaction is then calculated as the absolute value of the weighted average of the lag indices, using the normalized coefficients as weights^5^. To derive a single, representative “regulation lag” score for each TF, the method then computes the mean of all its associated interaction-specific lag scores across all of its TGs.

## Code availability

scMagnify is released under the BSD-3-Clause license, with code available at https://github.com/LiHongCSBLab/scMagnify. Code to reproduce the results in the paper can be found at https://github.com/LiHongCSBLab/scmagnify_reproducibility.

## Data availability

The human hematopoiesis, pancreatic endocrine and human kidney injury data presented and used in this study are publicly available via the original publications; We provide additional access to each of these datasets via a figshare collection: https://figshare.com/articles/dataset/scMagnify_reproducibility_datasets/30600797.

## Acknowledgements

This work was supported by Noncommunicable Chronic Diseases-National Science and Technology Major Project (2024ZD0531300), National Natural Science Foundation of China (32470707, 32300555) and Shanghai Municipal Science and Technology Major Project.

## Contributions

X.C. and H.L. contributed to the initial conceptualization and design of the project. H.L. supervised and conceived the project. X.C. designed and developed the scMagnify algorithms, performed the benchmark tasks and analyzed the datasets, and X.Y., B.S., H.W., and H.L. helped with the data analysis and discussed the results. X.Y., Z.T., and Y.Z., contributed to the software test. H.L., X.Y., P.L., H.Z., and Y.L. contributed to the writing review. X.C. and H.L. wrote the manuscript with help from other authors. All authors have reviewed and approved the final manuscript.

## References

1. Spitz, F. & Furlong, E. E. M. Transcription factors: from enhancer binding to developmental control. Nat Rev Genet 13, 613–626 (2012).

2. Lambert, S. A. et al. The Human Transcription Factors. Cell 172, 650–665 (2018).

3. Stadhouders, R., Filion, G. J. & Graf, T. Transcription factors and 3D genome conformation in cell-fate decisions. Nature 569, 345–354 (2019).

4. Schoenfelder, S. & Fraser, P. Long-range enhancer–promoter contacts in gene expression control. Nature Reviews Genetics 20, 437–455 (2019).

5. Kim, S. & Wysocka, J. Deciphering the multi-scale, quantitative cis-regulatory code. Molecular Cell 83, 373–392 (2023).

6. Mayor, R. Cell fate decisions during development. Science 364, 937–938 (2019).

7. Davidson, E. & Levin, M. Gene regulatory networks. Proceedings of the National Academy of Sciences 102, 4935–4935 (2005).

8. Karlebach, G. & Shamir, R. Modelling and analysis of gene regulatory networks. Nat Rev Mol Cell Biol 9, 770–780 (2008).

9. Langfelder, P. & Horvath, S. WGCNA: an R package for weighted correlation network analysis. BMC Bioinformatics 9, 559 (2008).

10. Huynh-Thu, V. A., Irrthum, A., Wehenkel, L. & Geurts, P. Inferring Regulatory Networks from Expression Data Using Tree-Based Methods. PLOS ONE 5, e12776 (2010).

11. Moerman, T. et al. GRNBoost2 and Arboreto: efficient and scalable inference of gene regulatory networks. Bioinformatics 35, 2159–2161 (2019).

12. Chan, T. E., Stumpf, M. P. H. & Babtie, A. C. Gene Regulatory Network Inference from Single-Cell Data Using Multivariate Information Measures. Cell Syst 5, 251–267.e3 (2017).

13. Banf, M. & Rhee, S. Y. Computational inference of gene regulatory networks: Approaches, limitations and opportunities. Biochimica et Biophysica Acta (BBA) - Gene Regulatory Mechanisms 1860, 41–52 (2017).

14. Pratapa, A., Jalihal, A. P., Law, J. N., Bharadwaj, A. & Murali, T. M. Benchmarking algorithms for gene regulatory network inference from single-cell transcriptomic data. Nat Methods 17, 147–154 (2020).

15. Matsumoto, H. et al. SCODE: an efficient regulatory network inference algorithm from single-cell RNA-Seq during differentiation. Bioinformatics 33, 2314–2321 (2017).

16. Papili Gao, N., Ud-Dean, S. M. M., Gandrillon, O. & Gunawan, R. SINCERITIES: inferring gene regulatory networks from time-stamped single cell transcriptional expression profiles. Bioinformatics 34, 258–266 (2018).

17. Qiu, X. et al. Inferring Causal Gene Regulatory Networks from Coupled Single-Cell Expression Dynamics Using Scribe. Cell Syst 10, 265–274.e11 (2020).

18. Wang, W., Wang, Y., Lyu, R. & Grün, D. Scalable identification of lineage-specific gene regulatory networks from metacells with NetID. Genome Biology 25, 275 (2024).

19. Singh, R., Wu, A. P., Mudide, A. & Berger, B. Causal gene regulatory analysis with RNA velocity reveals an interplay between slow and fast transcription factors. Cell Systems 15, 462–474.e5 (2024).

20. Boyle, A. P. et al. High-Resolution Mapping and Characterization of Open Chromatin across the Genome. Cell 132, 311–322 (2008).

21. Buenrostro, J. D., Giresi, P. G., Zaba, L. C., Chang, H. Y. & Greenleaf, W. J. Transposition of native chromatin for fast and sensitive epigenomic profiling of open chromatin, DNA-binding proteins and nucleosome position. Nat Methods 10, 1213–1218 (2013).

22. Johnson, D. S., Mortazavi, A., Myers, R. M. & Wold, B. Genome-wide mapping of in vivo protein-DNA interactions. Science 316, 1497–1502 (2007).

23. Kamal, A. et al. GRaNIE and GRaNPA: inference and evaluation of enhancer-mediated gene regulatory networks. Molecular Systems Biology n/a, e11627 (2023).

24. Klemm, S. L., Shipony, Z. & Greenleaf, W. J. Chromatin accessibility and the regulatory epigenome. Nat Rev Genet 20, 207–220 (2019).

25. Preissl, S., Gaulton, K. J. & Ren, B. Characterizing cis-regulatory elements using single-cell epigenomics. Nat Rev Genet 24, 21–43 (2023).

26. Badia-i-Mompel, P. et al. Gene regulatory network inference in the era of single-cell multi-omics. Nat Rev Genet 1–16 (2023) doi:10.1038/s41576-023-00618-5.

27. Aibar, S. et al. SCENIC: single-cell regulatory network inference and clustering. Nat Methods 14, 1083–1086 (2017).

28. Fleck, J. S. et al. Inferring and perturbing cell fate regulomes in human brain organoids. Nature 621, 365–372 (2023).

29. Kartha, V. K. et al. Functional inference of gene regulation using single-cell multi-omics. Cell Genomics 2, 100166 (2022).

30. Bravo González-Blas, C. et al. SCENIC+: single-cell multiomic inference of enhancers and gene regulatory networks. Nat Methods 1–13 (2023) doi:10.1038/s41592-023-01938-4.

31. Yuan, Q. & Duren, Z. Inferring gene regulatory networks from single-cell multiome data using atlas-scale external data. Nat Biotechnol 1–11 (2024) doi:10.1038/s41587-024-02182-7.

32. Kamimoto, K. et al. Dissecting cell identity via network inference and in silico gene perturbation. Nature 1–10 (2023) doi:10.1038/s41586-022-05688-9.

33. Wang, L. et al. Dictys: dynamic gene regulatory network dissects developmental continuum with single-cell multiomics. Nat Methods 1–11 (2023) doi:10.1038/s41592-023-01971-3.

34. Marcinkevičs, R. & Vogt, J. E. Interpretable Models for Granger Causality Using Self-explaining Neural Networks. Preprint at 10.48550/arXiv.2101.07600 (2021).

35. Fan, C., Wang, Y., Zhang, Y. & Ouyang, W. Interpretable Multi-Scale Neural Network for Granger Causality Discovery. in ICASSP 2023 - 2023 IEEE International Conference on Acoustics, Speech and Signal Processing (ICASSP) 1–5 (IEEE, Rhodes Island, Greece, 2023). doi:10.1109/ICASSP49357.2023.10096964.

36. Rabanser, S., Shchur, O. & Günnemann, S. Introduction to Tensor Decompositions and their Applications in Machine Learning. Preprint at 10.48550/arXiv.1711.10781 (2017).

37. Faure, L., Soldatov, R., Kharchenko, P. V. & Adameyko, I. scFates: a scalable python package for advanced pseudotime and bifurcation analysis from single-cell data. Bioinformatics 39, btac746 (2023).

38. Stormo, G. D. DNA binding sites: representation and discovery. Bioinformatics 16, 16–23 (2000).

39. Wu, A. P., Singh, R., Walsh, C. A. & Berger, B. Unveiling causal regulatory mechanisms through cell-state parallax. Nat Commun 16, 8096 (2025).

40. Badia-i-Mompel, P. et al. decoupleR: ensemble of computational methods to infer biological activities from omics data. Bioinformatics Advances 2, vbac016 (2022).

41. Morgunova, E. & Taipale, J. Structural perspective of cooperative transcription factor binding. Current Opinion in Structural Biology 47, 1–8 (2017).

42. Reiter, F., Wienerroither, S. & Stark, A. Combinatorial function of transcription factors and cofactors. Current Opinion in Genetics & Development 43, 73–81 (2017).

43. Browaeys, R., Saelens, W. & Saeys, Y. NicheNet: modeling intercellular communication by linking ligands to target genes. Nat Methods 17, 159–162 (2020).

44. Cheng, J., Zhang, J., Wu, Z. & Sun, X. Inferring microenvironmental regulation of gene expression from single-cell RNA sequencing data using scMLnet with an application to COVID-19. Brief Bioinform 22, 988–1005 (2021).

45. He, C., Zhou, P. & Nie, Q. exFINDER: identify external communication signals using single-cell transcriptomics data. Nucleic Acids Research 51, e58–e58 (2023).

46. Li, D. et al. TraSig: inferring cell-cell interactions from pseudotime ordering of scRNA-Seq data. Genome Biology 23, 73 (2022).

47. Li, H., Zhang, Z., Squires, M., Chen, X. & Zhang, X. scMultiSim: simulation of single-cell multi-omics and spatial data guided by gene regulatory networks and cell–cell interactions. Nat Methods 1–12 (2025) doi:10.1038/s41592-025-02651-0.

48. Taing, L. et al. Cistrome Data Browser: integrated search, analysis and visualization of chromatin data. Nucleic Acids Research 52, D61–D66 (2024).

49. Hammelman, J., Patel, T., Closser, M., Wichterle, H. & Gifford, D. Ranking reprogramming factors for cell differentiation. Nat Methods 19, 812–822 (2022).

50. Weiler, P., Lange, M., Klein, M., Pe’er, D. & Theis, F. CellRank 2: unified fate mapping in multiview single-cell data. Nat Methods 1–10 (2024) doi:10.1038/s41592-024-02303-9.

51. Persad, S. et al. SEACells infers transcriptional and epigenomic cellular states from single-cell genomics data. Nat Biotechnol 1–12 (2023) doi:10.1038/s41587-023-01716-9.

52. Orkin, S. H. & Zon, L. I. Hematopoiesis: an evolving paradigm for stem cell biology. Cell 132, 631–644 (2008).

53. Pongubala, J. M. R. et al. Transcription factor EBF restricts alternative lineage options and promotes B cell fate commitment independently of Pax5. Nat Immunol 9, 203–215 (2008).

54. Acquaviva, J., Chen, X. & Ren, R. IRF-4 functions as a tumor suppressor in early B-cell development. Blood 112, 3798–3806 (2008).

55. Mansson, R. et al. Positive intergenic feedback circuitry, involving EBF1 and FOXO1, orchestrates B-cell fate. Proceedings of the National Academy of Sciences 109, 21028–21033 (2012).

56. Nechanitzky, R. et al. Transcription factor EBF1 is essential for the maintenance of B cell identity and prevention of alternative fates in committed cells. Nat Immunol 14, 867–875 (2013).

57. Peters, I. J. A., de Pater, E. & Zhang, W. The role of GATA2 in adult hematopoiesis and cell fate determination. Front. Cell Dev. Biol. 11, (2023).

58. Siatecka, M. & Bieker, J. J. The multifunctional role of EKLF/KLF1 during erythropoiesis. Blood 118, 2044–2054 (2011).

59. Friedman, A. D. Transcriptional control of granulocyte and monocyte development. Oncogene 26, 6816–6828 (2007).

60. Loughran, S. J. et al. The transcription factor Erg is essential for definitive hematopoiesis and the function of adult hematopoietic stem cells. Nat Immunol 9, 810–819 (2008).

61. Itoh-Nakadai, A. et al. A Bach2-Cebp Gene Regulatory Network for the Commitment of Multipotent Hematopoietic Progenitors. Cell Reports 18, 2401–2414 (2017).

62. Johanson, T. M. et al. Transcription-factor-mediated supervision of global genome architecture maintains B cell identity. Nat Immunol 19, 1257–1264 (2018).

63. Chen, Y. et al. GraphVelo allows for accurate inference of multimodal velocities and molecular mechanisms for single cells. Nat Commun 16, 7831 (2025).

64. Lin, Y. C. et al. A global network of transcription factors, involving E2A, EBF1 and Foxo1, that orchestrates B cell fate. Nat Immunol 11, 635–643 (2010).

65. Hu, Y. et al. Multiscale footprints reveal the organization of cis-regulatory elements. Nature 1–8 (2025) doi:10.1038/s41586-024-08443-4.

66. Chen, C.-H. et al. Determinants of transcription factor regulatory range. Nat Commun 11, 2472 (2020).

67. Fischer, U. et al. Cell Fate Decisions: The Role of Transcription Factors in Early B-cell Development and Leukemia. Blood Cancer Discovery 1, 224–233 (2020).

68. Somasundaram, R., Prasad, M. A. J., Ungerbäck, J. & Sigvardsson, M. Transcription factor networks in B-cell differentiation link development to acute lymphoid leukemia. Blood 126, 144–152 (2015).

69. Schwickert, T. A. et al. Stage-specific control of early B cell development by the transcription factor Ikaros. Nat Immunol 15, 283–293 (2014).

70. Ficara, F., Murphy, M. J., Lin, M. & Cleary, M. L. Pbx1 Regulates Self-Renewal of Long-Term Hematopoietic Stem Cells by Maintaining Their Quiescence. Cell Stem Cell 2, 484–496 (2008).

71. Doulatov, S. et al. Induction of Multipotential Hematopoietic Progenitors from Human Pluripotent Stem Cells via Respecification of Lineage-Restricted Precursors. Cell Stem Cell 13, 459–470 (2013).

72. Voit, R. A. & Sankaran, V. G. MECOM Deficiency: from Bone Marrow Failure to Impaired B-Cell Development. J Clin Immunol 43, 1052–1066 (2023).

73. Huang, J., Wang, Y., Liu, J., Chu, M. & Wang, Y. TFDP3 as E2F Unique Partner, Has Crucial Roles in Cancer Cells and Testis. Front. Oncol. 11, (2021).

74. Ishii, S. et al. Genome-wide ATAC-see screening identifies TFDP1 as a modulator of global chromatin accessibility. Nat Genet 1–10 (2024) doi:10.1038/s41588-024-01658-1.

75. Ng, A. P. et al. An Erg-driven transcriptional program controls B cell lymphopoiesis. Nat Commun 11, 3013 (2020).

76. Sam-Yellowe, T. Y. B Cell Development, Activation and Immunoglobulin Structure. in Immunology: Overview and Laboratory Manual (ed. Sam-Yellowe, T. Y.) 73–80 (Springer International Publishing, Cham, 2021). doi:10.1007/978-3-030-64686-8_10.

77. Azagra, A. et al. In vivo conditional deletion of HDAC7 reveals its requirement to establish proper B lymphocyte identity and development. J Exp Med 213, 2591–2601 (2016).

78. Zhang, X. et al. Pax6 is regulated by Meis and Pbx homeoproteins during pancreatic development. Developmental Biology 300, 748–757 (2006).

79. Kobberup, S., Nyeng, P., Juhl, K., Hutton, J. & Jensen, J. ETS-family genes in pancreatic development. Developmental Dynamics 236, 3100–3110 (2007).

80. Gage, B. K. et al. The Role of ARX in Human Pancreatic Endocrine Specification. PLOS ONE 10, e0144100 (2015).

81. Hang, Y. & Stein, R. MafA and MafB activity in pancreatic β cells. Trends in Endocrinology & Metabolism 22, 364–373 (2011).

82. Gradwohl, G., Dierich, A., LeMeur, M. & Guillemot, F. neurogenin3 is required for the development of the four endocrine cell lineages of the pancreas. Proceedings of the National Academy of Sciences 97, 1607–1611 (2000).

83. Wilson, M. E. et al. The HMG Box Transcription Factor Sox4 Contributes to the Development of the Endocrine Pancreas. Diabetes 54, 3402–3409 (2005).

84. Soyer, J. et al. Rfx6 is an Ngn3-dependent winged helix transcription factor required for pancreatic islet cell development. Development 137, 203–212 (2010).

85. Xu, E. E. et al. SOX4 cooperates with neurogenin 3 to regulate endocrine pancreas formation in mouse models. Diabetologia 58, 1013–1023 (2015).

86. Hobert, O. Regulatory logic of neuronal diversity: Terminal selector genes and selector motifs. Proceedings of the National Academy of Sciences 105, 20067–20071 (2008).

87. Collombat, P. et al. Opposing actions of Arx and Pax4 in endocrine pancreas development. Genes Dev. 17, 2591–2603 (2003).

88. Zhang, C. et al. MafA is a key regulator of glucose-stimulated insulin secretion. Mol Cell Biol 25, 4969–4976 (2005).

89. Gittes, G. K. Developmental biology of the pancreas: A comprehensive review. Developmental Biology 326, 4–35 (2009).

90. Armingol, E., Baghdassarian, H. M. & Lewis, N. E. The diversification of methods for studying cell–cell interactions and communication. Nat Rev Genet 1–20 (2024) doi:10.1038/s41576-023-00685-8.

91. Gisch, D. L. et al. The chromatin landscape of healthy and injured cell types in the human kidney. Nat Commun 15, 433 (2024).

92. Li, D. et al. Role of promoting inflammation of Krüppel-like factor 6 in acute kidney injury. Renal Failure 42, 693–703 (2020).

93. Sun, D., Liu, X., Zhu, L. & Zhang, B. Zinc-finger E-box-binding homeobox 1 alleviates acute kidney injury by activating autophagy and the AMPK/mTOR pathway. Molecular Medicine Reports 23, 1–9 (2021).

94. Lake, B. B. et al. An atlas of healthy and injured cell states and niches in the human kidney. Nature 619, 585–594 (2023).

95. Reck, M. et al. Multiomic analysis of human kidney disease identifies a tractable inflammatory and pro-fibrotic tubular cell phenotype. Nat Commun 16, 4745 (2025).

96. Ma, J. et al. CD44 Is a Prognostic Biomarker Correlated With Immune Infiltrates and Metastasis in Clear Cell Renal Cell Carcinoma. Anticancer Research 43, 3493–3506 (2023).

97. La Manno, G. et al. RNA velocity of single cells. Nature 560, 494–498 (2018).

98. Bergen, V., Soldatov, R. A., Kharchenko, P. V. & Theis, F. J. RNA velocity—current challenges and future perspectives. Molecular Systems Biology 17, e10282 (2021).

99. Wang, W. et al. RegVelo: gene-regulatory-informed dynamics of single cells. 2024.12.11.627935 Preprint at 10.1101/2024.12.11.627935 (2024).

100. Lederer, A. R. et al. Statistical inference with a manifold-constrained RNA velocity model uncovers cell cycle speed modulations. Nat Methods 21, 2271–2286 (2024).

101. Furtwängler, B. et al. Mapping early human blood cell differentiation using single-cell proteomics and transcriptomics. Science https://www.science.org/doi/10.1126/science.adr8785 (2025).

102. Chen, M., Fu, R., Chen, Y., Li, L. & Wang, S.-W. High-resolution, noninvasive single-cell lineage tracing in mice and humans based on DNA methylation epimutations. Nat Methods 22, 488–498 (2025).

103. Bunne, C. et al. How to build the virtual cell with artificial intelligence: Priorities and opportunities. Cell 187, 7045–7063 (2024).

104. Noutahi, E. et al. Virtual Cells: Predict, Explain, Discover. Preprint at 10.48550/arXiv.2505.14613 (2025).

105. Ma, S. et al. Chromatin Potential Identified by Shared Single-Cell Profiling of RNA and Chromatin. Cell 183, 1103–1116.e20 (2020).

106. Korhonen, J., Martinmäki, P., Pizzi, C., Rastas, P. & Ukkonen, E. MOODS: fast search for position weight matrix matches in DNA sequences. https://doi.org/10.1093/bioinformatics/btp554 doi:10.1093/bioinformatics/btp554.

107. Kulakovskiy, I. V. et al. HOCOMOCO: towards a complete collection of transcription factor binding models for human and mouse via large-scale ChIP-Seq analysis. https://doi.org/10.1093/nar/gkx1106 doi:10.1093/nar/gkx1106.

108. Sims, C. A. Macroeconomics and Reality. Econometrica 48, 1 (1980).

109. Yuan, M. & Lin, Y. Model Selection and Estimation in Regression with Grouped Variables. https://doi.org/10.1111/j.1467-9868.2005.00532.x doi:10.1111/j.1467-9868.2005.00532.x.

110. Zitnik, M. et al. Current and future directions in network biology. Preprint at 10.48550/arXiv.2309.08478 (2023).

111. Otto, D. J., Jordan, C., Dury, B., Dien, C. & Setty, M. Quantifying cell-state densities in single-cell phenotypic landscapes using Mellon. Nat Methods 1–11 (2024) doi:10.1038/s41592-024-02302-w.

112. Klein, D. et al. Mapping cells through time and space with moscot. Nature 1–11 (2025) doi:10.1038/s41586-024-08453-2.

113. Ibarra, I. L. et al. Learning sequence-based regulatory dynamics in single-cell genomics. 2024.08.07.605876 Preprint at 10.1101/2024.08.07.605876 (2024).

114. Haghverdi, L., Büttner, M., Wolf, F. A., Buettner, F. & Theis, F. J. Diffusion pseudotime robustly reconstructs lineage branching. Nat Methods 13, 845–848 (2016).

115. Wolf, F. A., Angerer, P. & Theis, F. J. SCANPY: large-scale single-cell gene expression data analysis. Genome Biology 19, 15 (2018).

116. Bravo González-Blas, C. et al. cisTopic: cis-regulatory topic modeling on single-cell ATAC-seq data. Nat Methods 16, 397–400 (2019).

