## Supplemental Figures for "Decomposing multi-scale dynamic regulation from single-cell multiomics with scMagnify"

### Supplementary Data

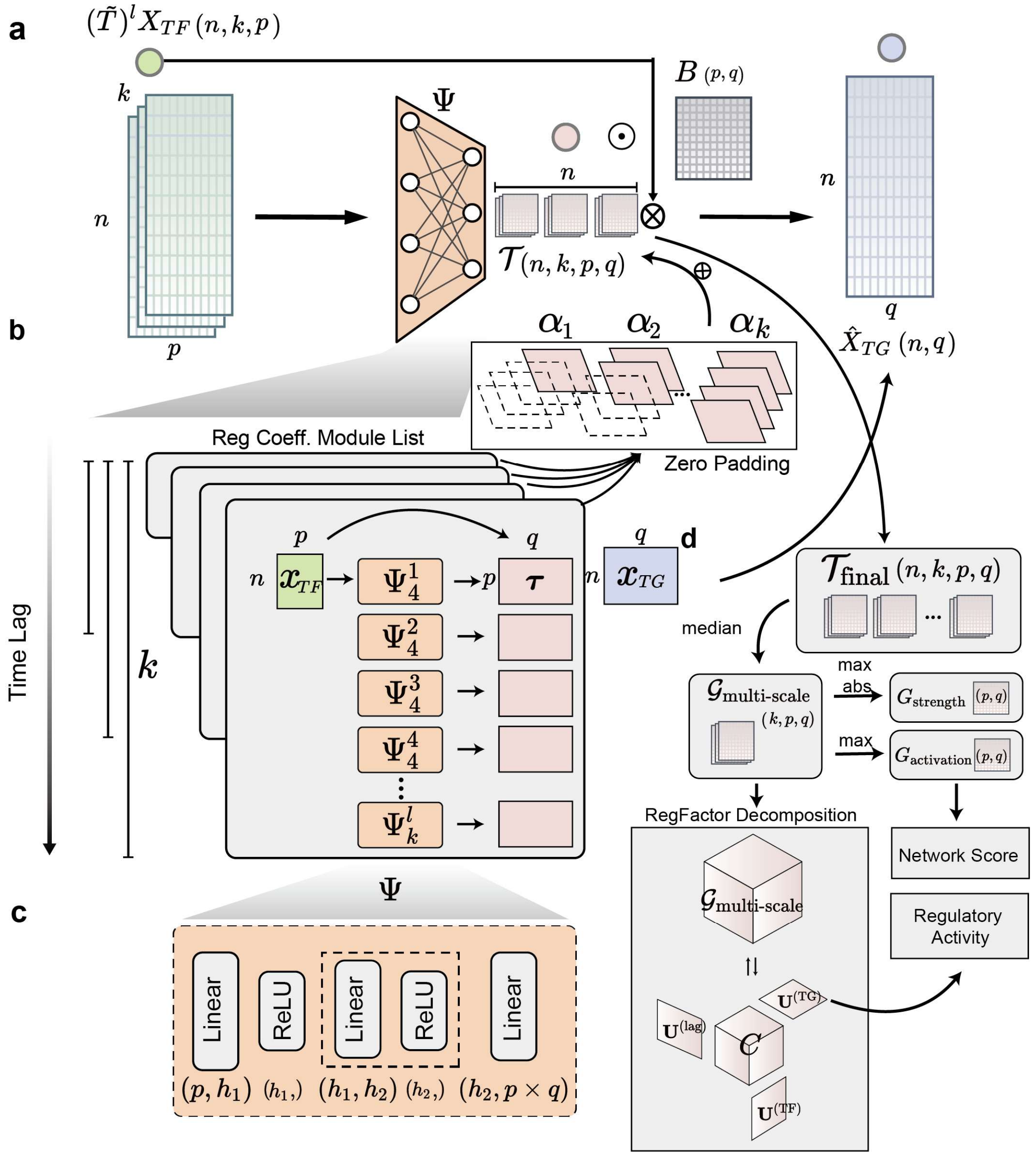


**Supplementary Fig. 1 | Detailed architecture of the scMagnify framework.**

**a,** The multi-scale self-explaining neural network model predicts target gene (TG) expression ($\hat{X}_{TG}$) from time-lagged and graph-diffused transcription factor (TF) expression ($(\tilde{T})^l X_{TF}$). The input is processed by a neural network ($\Psi$) to generate regulatory coefficients ($\mathcal{T}$), which are constrained by a TF binding network ($B$). An attention mechanism ($\alpha_k$) learns to weight and integrate signals from each time-lag branch.

**b,** The model uses a parallel-branch architecture to capture regulatory effects across multiple time lags ($k$). Each branch $k$ aggregates information from sub-networks ($\Psi_k^l$) representing $l$ historical steps.

**c,** Each $\Psi$ sub-module is a feedforward neural network with ReLU activations that maps TF inputs to a TF-TG regulatory coefficient matrix.

**d,** Model outputs ($\mathcal{T}_{\text{final}}$) are aggregated via median to build a robust multi-scale GRN ($\mathcal{G}_{\text{multi-scale}}$).

This tensor is used to: (1) derive ensemble networks ($G_{strength}$ and $G_{activation}$); and (2) perform RegFactor Decomposition via Tucker decomposition to identify regulatory modules (core tensor $C$ and factor matrices $\mathbf{U}$). These results are used for downstream Network Score and Regulatory Activity analyses. S

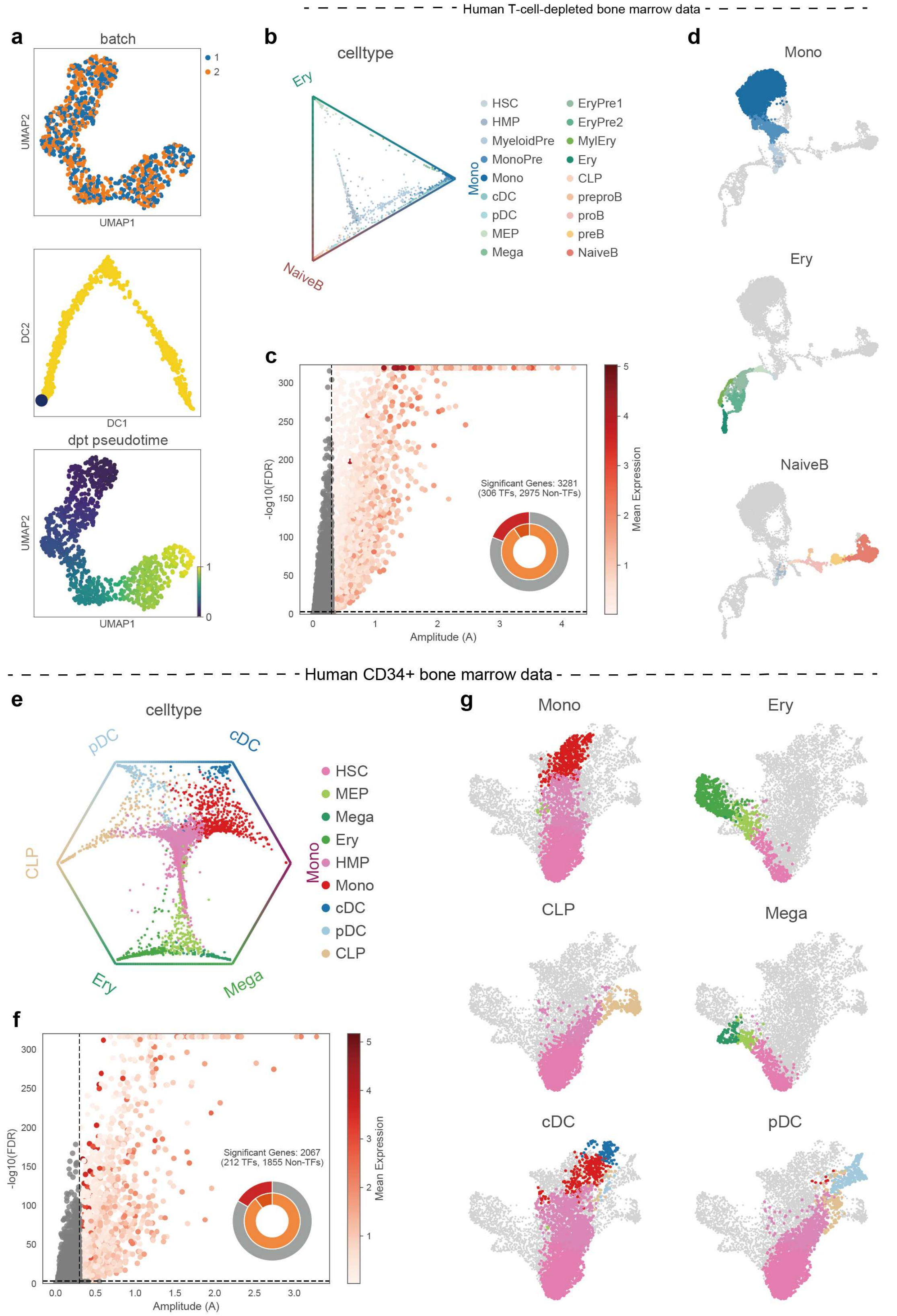

**Supplementary Fig. 2 | Preprocessing of synthetic and real-world datasets for benchmarking.**
**a,** Preprocessing of the synthetic dataset generated by scMultiSim. UMAP embedding of simulated cells colored by batch (top), diffusion map embedding and root cell for pseudotime analysis (middle), and UMAP colored by the inferred dpt pseudotime (bottom).
**b-d,** Preprocessing of the human T-cell-depleted bone marrow dataset. **b,** Ternary plot showing cell fate probabilities towards the Ery, Mono, and NaiveB lineages, colored by celltype. **c,** Volcano plot identifying 3,281 genes significantly associated with pseudotime (defined by Amplitude (A) > 0.3 and FDR < 0.001), plotting Amplitude (A) against statistical significance (-log10(FDR)). The donut chart summarizes the significant genes (306 TFs and 2975 non-TFs). **d,** UMAP plots highlighting cells assigned to the terminal Mono, Ery, and NaiveB lineages.
**e-g,** Preprocessing of the human CD34+ bone marrow dataset. **e,** Ternary plot showing cell fate probabilities across six major lineages (CLP, pDC, cDC, Mono, Mega, Ery), colored by celltype.
**f,** Volcano plot identifying 2,067 significant genes associated with pseudotime (defined by Amplitude (A) > 0.3 and FDR < 0.001), plotting Amplitude (A) against statistical significance (-log10(FDR)). The donut chart summarizes the significant genes (212 TFs and 1855 non-TFs). **g,** UMAP plots highlighting cells assigned to the terminal Mono, Ery, CLP, Mega, cDC, and pDC lineages.

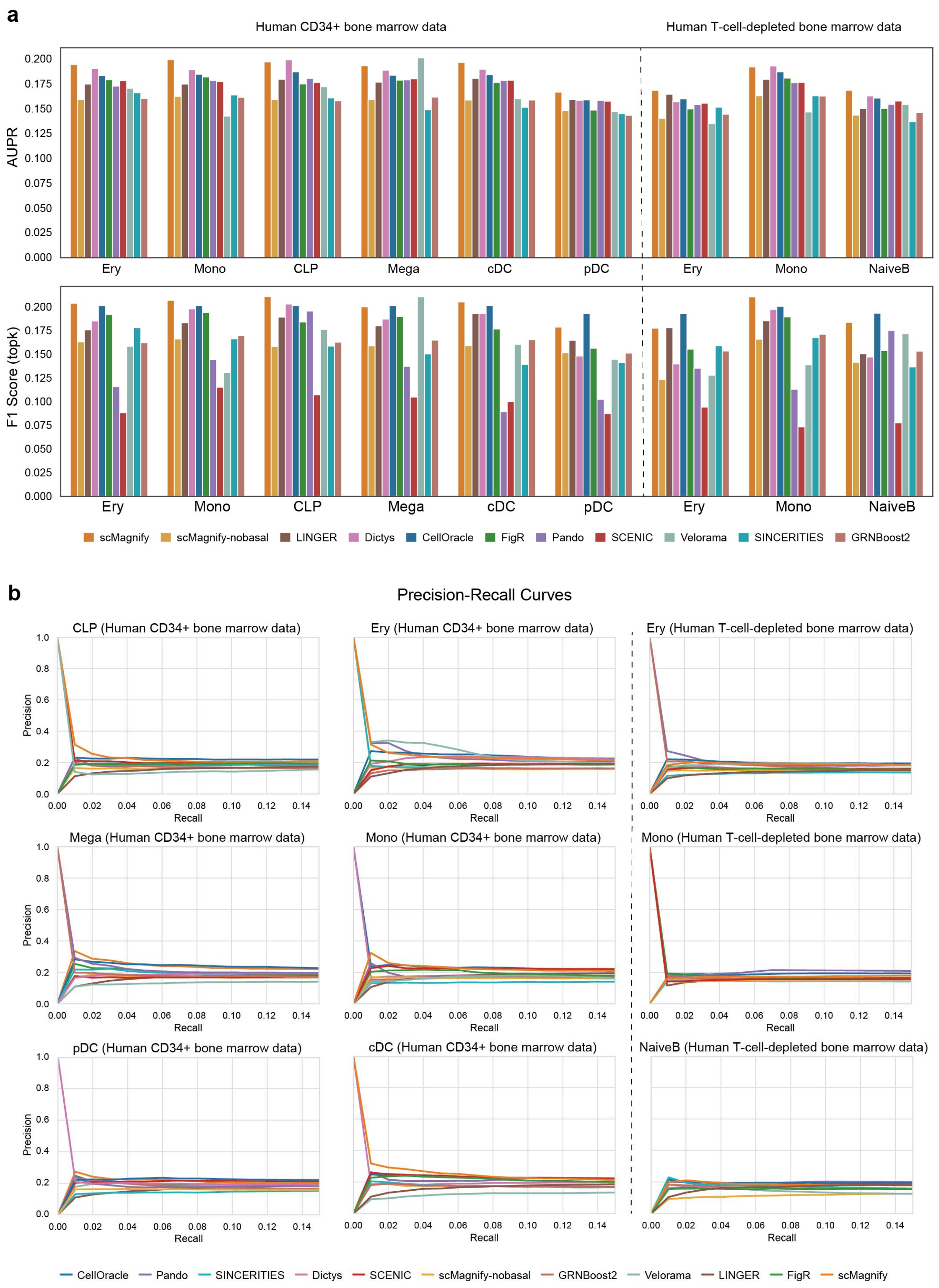


**Supplementary Fig. 3 | Detailed performance of TF-TG interaction accuracy.**

**a,** Bar plots comparing AUPR (top panel) and F1-Score (bottom panel) for scMagnify and all competing methods across the nine tested lineages from the two bone marrow datasets. These plots show the full data for each lineage, which is presented as a distribution in the main **Fig. 2c**.

**b,** Full Precision-Recall (PR) curves for all benchmarked methods, shown for each of the nine individual lineages. The Area Under the PR curve (AUPR) is calculated from these curves.

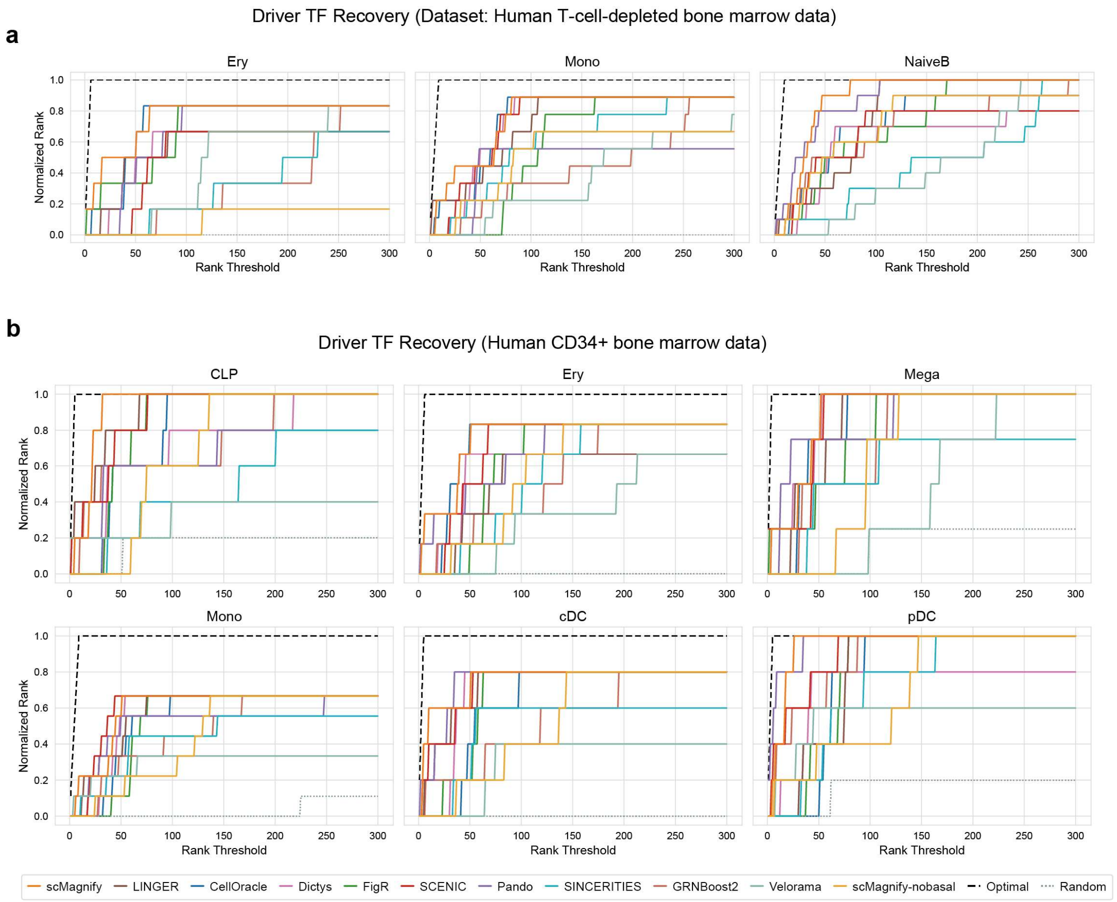


**Supplementary Fig. 4 | Detailed rank recovery curves for the driver TF benchmark.**

**a,** Rank recovery curves for the three lineages (Ery, Mono, NaiveB) from the human T-cell-depleted bone marrow dataset. **b,** Rank recovery curves for the six lineages (CLP, Ery, Mega, Mono, cDC, pDC) from the human CD34+ bone marrow dataset.

For each plot, the Normalized Rank (y-axis) shows the fraction of known driver TFs recovered within the top Rank Threshold (x-axis) of TFs, as ranked by degree centrality. These curves represent the detailed data summarized in the main **Fig. 2e**.

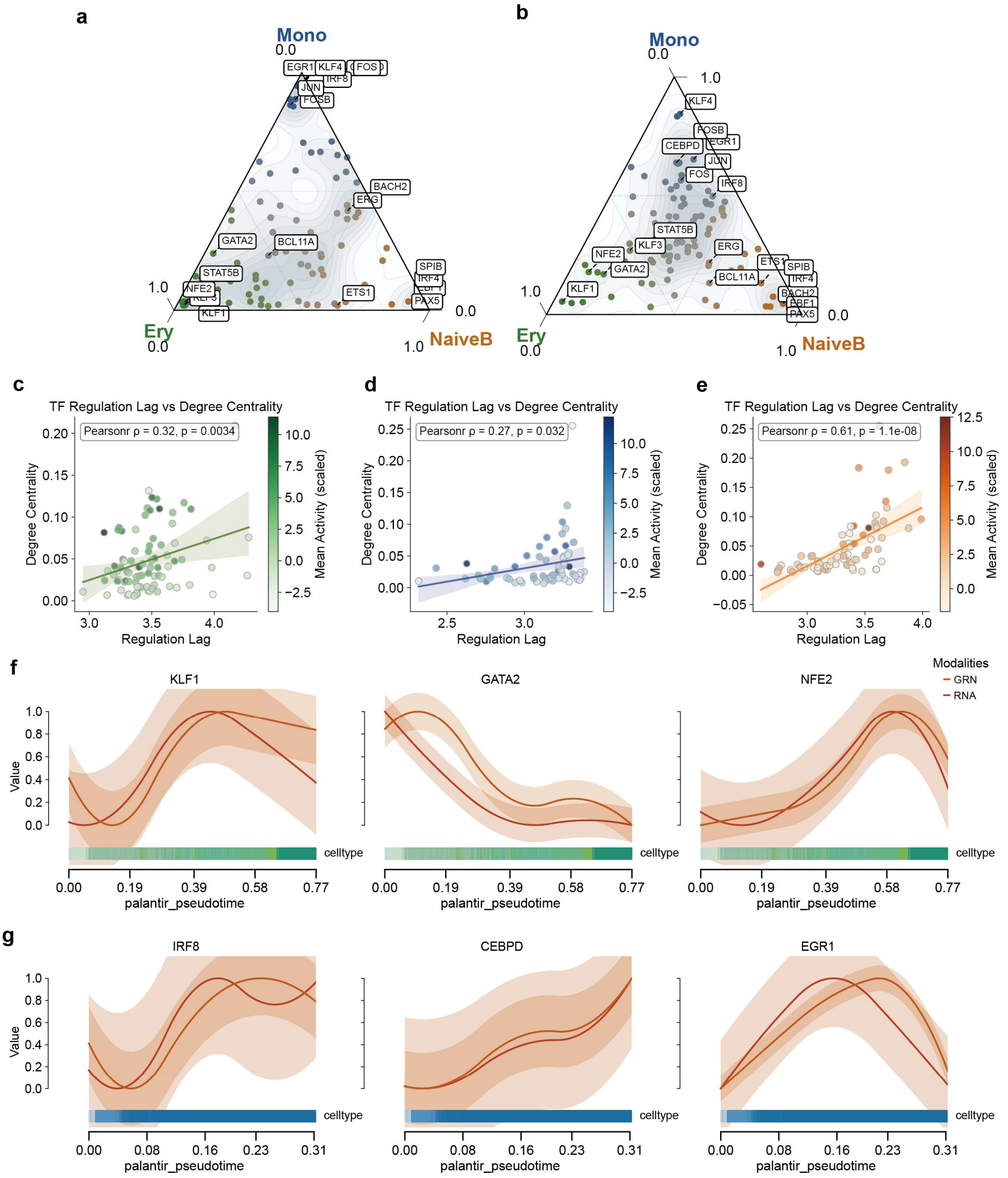


**Supplementary Fig. 5 | Regulation specificity and regulation lag analysis in human hematopoiesis.**

**a, b,** Ternary plots showing the lineage specificity of TFs based on their mean regulatory activity (**a**) and mean RNA expression (**b**). TFs are positioned according to their specificity towards the Ery, Mono, and NaiveB lineages. These plots supplement the analysis in the main **Fig. 3b**.

**c, d, e,** Scatter plots showing the positive correlation between Regulation Lag and Degree Centrality for TFs in the Ery (**c**), Mono (**d**), and NaiveB (**e**) lineages. Points are colored by their mean regulatory activity. Pearson correlation coefficients ( ρ ) and p-values are shown.

**f, g,** Line plots showing the temporal dynamics of normalized RNA expression (orange line) and inferred regulatory activity (purple line) along the pseudotime axis for key Erythroid TFs (**f**) and

Monocyte TFs (**g**).

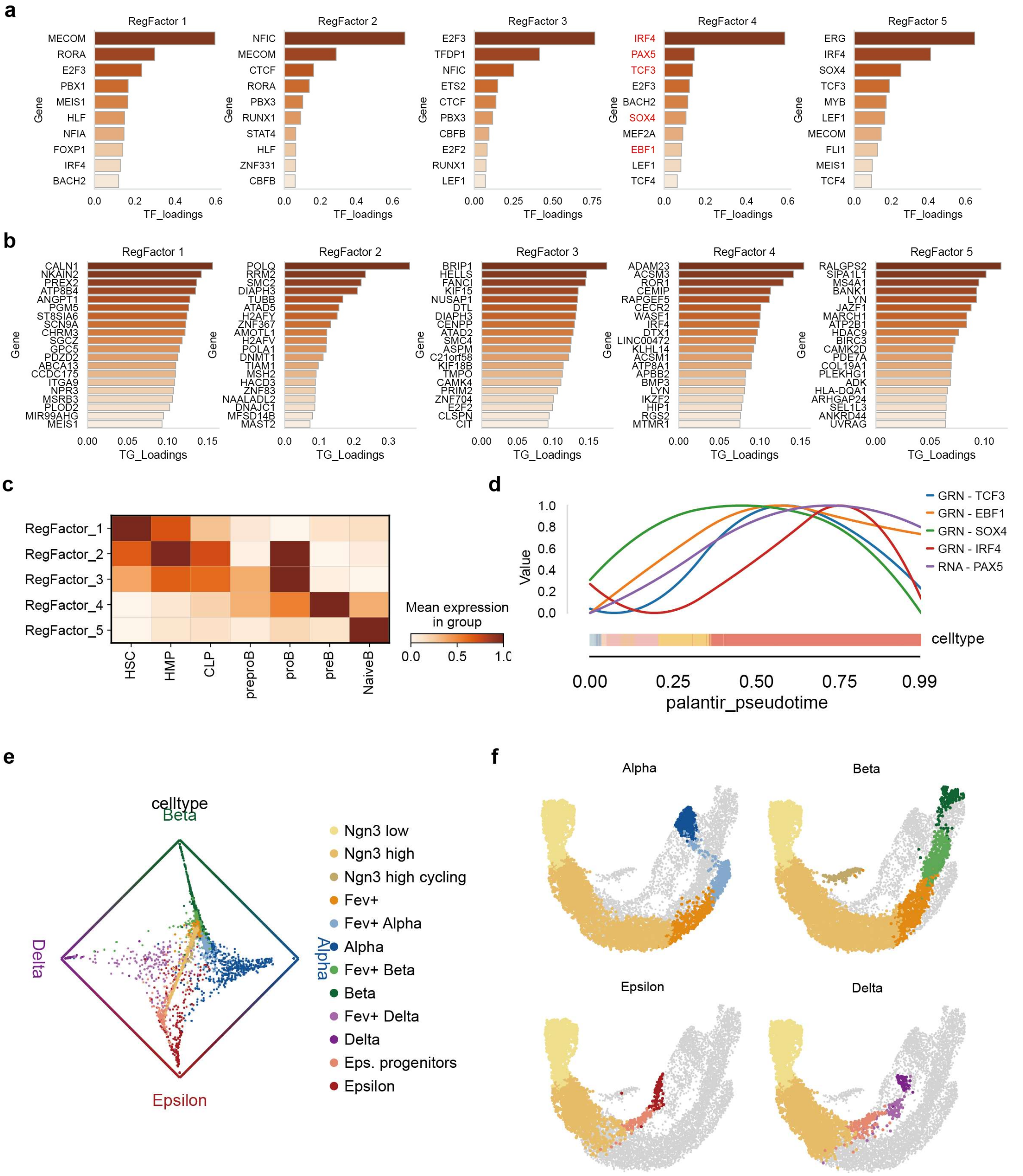


**Supplementary Fig. 6 | Additional results for B cell commitment and pancreas development.**

**a, b,** Bar plots showing the top 10 TF loadings (**a**) and TG loadings (**b**) for each of the 5 RegFactors identified in the Naive B-cell lineage, related to main **Fig. 4a-b**.

**c,** Violin plots displaying the inferred regulatory activity of each of the 5 B-cell RegFactors across the B-cell differentiation trajectory. This supplements the UMAP projections in main **Fig. 4b**.

**d,** Line plots showing the temporal dynamics of the inferred regulatory activity for the core RegFactor 4 TFs (TCF3, EBF1, SOX4, IRF4) and the RNA expression of their target *PAX5*, plotted along the B-cell lineage pseudotime. This is related to the main **Fig. 4d-f**.

**e,** Ternary plot showing cell fate probabilities for the mouse embryonic pancreas dataset, colored by cell type.

**f,** UMAP plots for the mouse pancreas dataset, highlighting the cells assigned to the Alpha, Beta, Delta, and Epsilon terminal lineages.

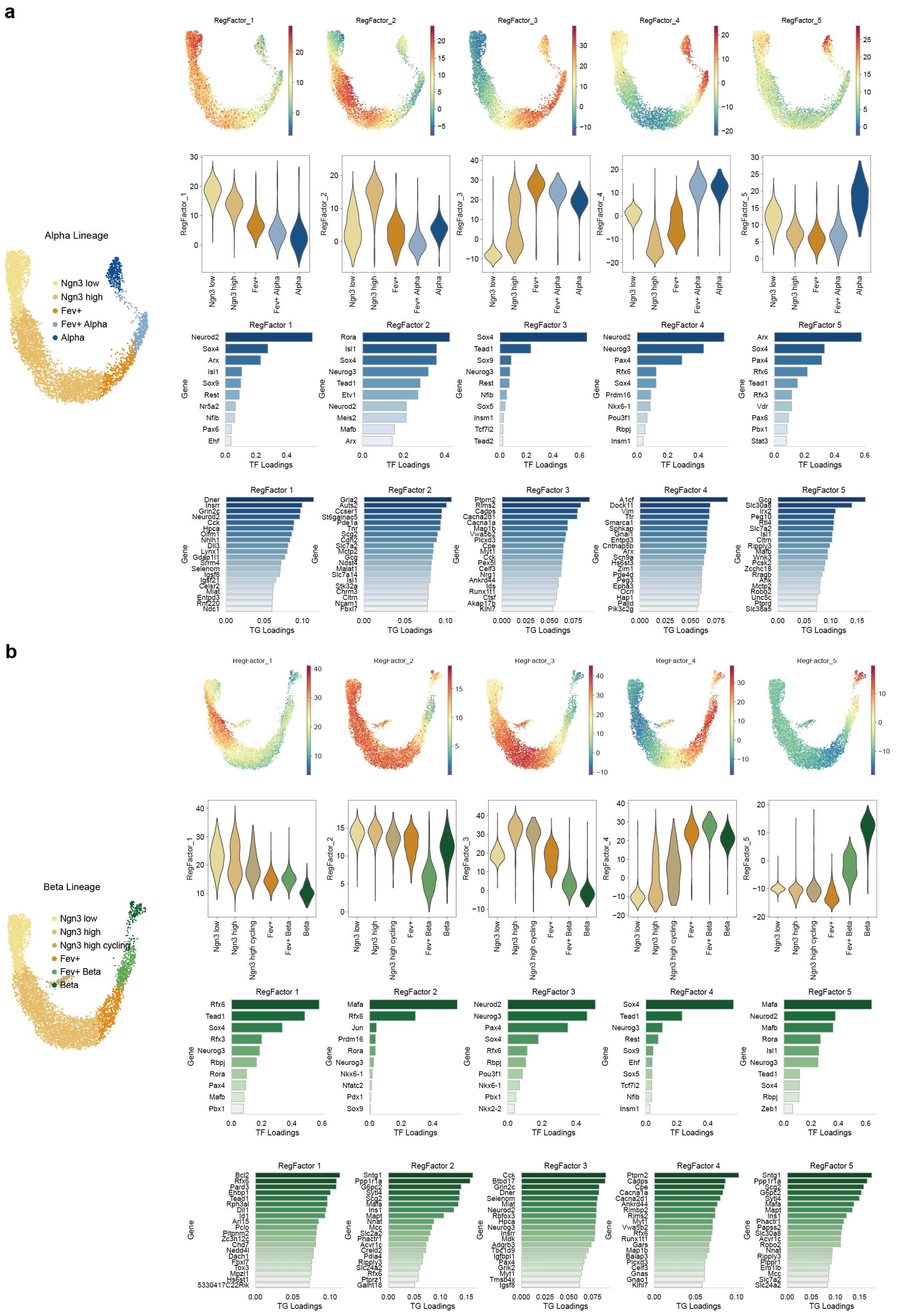


**Supplementary Fig. 7 | Characterization of RegFactors in mouse pancreas alpha and beta cell differentiation.**

This figure details the RegFactors identified for both the Alpha and Beta lineages. For each lineage, the leftmost UMAP shows the cells included in the trajectory. For each of the 5 RegFactors, the plots show: (top) UMAP projection of the RegFactor's activity; (middle) Violin plot of the RegFactor's activity across cell types in the differentiation path; (bottom left) Bar plot of the top TF loadings; (bottom right) Bar plot of the top TG loadings. **a,** Analysis of the 5 RegFactors for the Alpha Lineage. **b,** Analysis of the 5 RegFactors for the Beta Lineage.

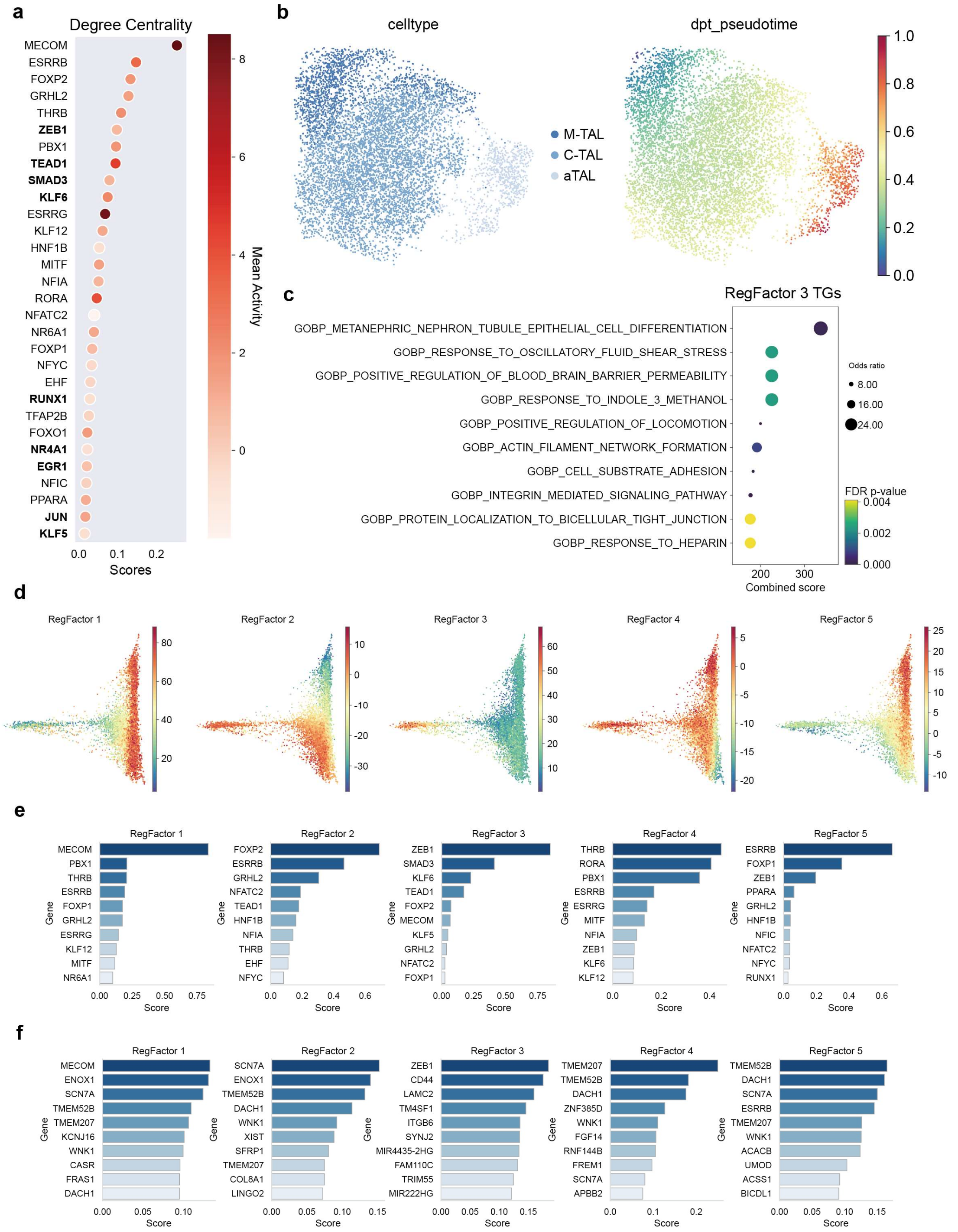


**Supplementary Fig. 8 | Additional results of the kidney injury dataset.**

**a,** Lollipop plot ranking TFs in the Thick Ascending Limb (TAL) lineage. The x-axis and dot size represent Degree Centrality, while the color indicates mean TF activity.

**b,** UMAP embedding of the TAL epithelial populations, colored by celltype (left panel; M-TAL, C-TAL, aTAL) and by dpt pseudotime (right panel), illustrating the injury trajectory. This corresponds to the main **Fig. 6b-c**.

**c,** Dot plot showing functional enrichment analysis (GO:BP terms) for the Top target genes (TGs) of RegFactor 3. Dot size represents the Odds ratio, and color represents the FDR p-value.

**d,** Diffusion Map projections showing the inferred activity of each of the 5 RegFactors identified in the TAL lineage.

**e, f,** Bar plots showing the top TF loadings (**e**) and top TG loadings (**f**) for each of the 5 TAL RegFactors. These plots provide the full data for all modules, supplementing the analysis of RegFactor 3 shown in the main **Fig. 6f**.

**Supplementary Tables 1. Comparison of scMagnify with existing single-cell multi-omic GRN inference methods.**

**Supplementary Tables 2. Curated list of gold-standard lineage-driving TFs used for benchmarking and analysis.**

**Supplementary Tables 3-6 Detailed RegFactor decomposition results.** These tables provide the complete quantitative results of the RegFactors identified by Tucker decomposition of the multi-scale GRNs for each biological lineage. The tables include the contribution weights (loadings) of transcription factors (TFs) for each RegFactor, as well as the target gene (TG) loadings and the time-lag loadings that define each RegFactor's activation profile.

Supplementary Table 3. RegFactor decomposition results for the Human T-cell-depleted bone marrow NaïveB-cell lineage.

Supplementary Table 4. RegFactor decomposition results for the Mouse Pancreas Alpha-cell lineage.

Supplementary Table 5. RegFactor decomposition results for the Mouse Pancreas Beta-cell lineage.

Supplementary Table 6. RegFactor decomposition results for the Human Kidney Injury TAL lineage.
